# Evolution of *E. coli* in a mouse model of inflammatory bowel disease leads to a disease-specific bacterial genotype and trade-offs with clinical relevance

**DOI:** 10.1101/2023.08.16.553450

**Authors:** Rahul Unni, Nadia Andrea Andreani, Marie Vallier, Silke S. Heinzmann, Jan Taubenheim, Martina A. Guggeis, Florian Tran, Olga Vogler, Sven Künzel, Jan-Bernd Hövener, Philip Rosenstiel, Christoph Kaleta, Astrid Dempfle, Daniel Unterweger, John F. Baines

## Abstract

Inflammatory bowel disease (IBD) is a persistent inflammatory condition that affects the gastrointestinal tract and presents significant challenges in its management and treatment. Despite the knowledge that within-host bacterial evolution occurs in the intestine, the disease has rarely been studied from an evolutionary perspective. In this study, we aimed to investigate the evolution of resident bacteria during intestinal inflammation and whether- and how disease-related bacterial genetic changes may present trade-offs with potential therapeutic importance. Here, we perform an *in vivo* evolution experiment of *E. coli* in a gnotobiotic mouse model of IBD, followed by multiomic analyses to identify disease-specific genetic and phenotypic changes in bacteria that evolved in an inflamed versus a non-inflamed control environment. Our results demonstrate distinct evolutionary changes in *E. coli* specific to inflammation, including a single nucleotide variant that independently reached high frequency in all inflamed mice. Using *ex vivo* fitness assays, we find that these changes are associated with a higher fitness in an inflamed environment compared to isolates derived from non-inflamed mice. Further, using large-scale phenotypic assays, we show that bacterial adaptation to inflammation results in clinically relevant phenotypes, which intriguingly include collateral sensitivity to antibiotics. Bacterial evolution in an inflamed gut yields specific genetic and phenotypic signatures. These results may serve as a basis for developing novel evolution-informed treatment approaches for patients with intestinal inflammation.

## Introduction

Dysbiosis of the intestinal microbial community, characterized by alterations in bacterial composition and function, is a hallmark of inflammatory bowel disease (IBD). The nature of these microbial imbalances has been described extensively and at numerous complementary levels, including diversity parameters, taxonomic- and functional genomic changes, and at the level of gene expression ^1–6^. This important body of work accounts for changes observed at the ecological level, while the latter also considers the potential contribution of phenotypic plasticity of gut microbes to disease susceptibility. Moreover, numerous studies have highlighted the association of dysbiotic microbial signatures with distinct clinical IBD subtypes, including Crohn’s disease (CD) and ulcerative colitis (UC), further emphasizing the intricate relationship between the gut microbiota and disease pathogenesis ^3,4,7^.

While research has focused extensively on the altered ecology of the gut microbiota in IBD, its potential for evolutionary change during disease pathogenesis has received comparatively limited attention. However, numerous recent studies have revealed the capacity of bacteria to undergo adaptive evolution within the host environment, including the gut [reviewed in ^8^]. This phenomenon is evident in both mouse models ^9–11^ and human subjects ^12^, even in the absence of overt disease. Importantly, recent studies indicate that evolutionary changes also occur in the context of host inflammation. Elhenawy et al. focused on CD-related adherent-invasive *E. coli* (AIEC) using a murine model of chronic colonization, and revealed the evolution of lineages displaying enhanced invasive and metabolic capabilities ^13^. Notably, they found that the fitness benefits conferred by increased motility were specific to the host environment, suggesting an evolutionary trade-off. In a second study exploring bacterial evolution in aging mice, significant differences in *E. coli* evolution between old and young mice were revealed ^14^. The aged mouse environment exhibited increased inflammation, leading to the specific targeting of stress-related functions in *E. coli*.

These findings emphasize the importance of including evolutionary perspectives when studying dysbiosis of the gut microbiome. In particular, documenting bacterial evolution within the inflamed gut has the potential to reveal the *timing* of disease-specific signatures, i.e., it may help disentangle the classical “chicken or egg” dilemma in microbiome research ^15^, as well as to shed light on potential trade-offs resulting from adaptive changes. Evolutionary trade-offs occur when an increase in fitness in one environment is accompanied by a decrease in fitness in another ^16,17^. These trade-offs can have significant therapeutic implications and are observed in various related fields, including antibiotic resistance ^18–21^ and cancer treatment ^22–25^. Thus, elucidating trade-offs associated with bacterial adaptations in the inflamed gut may offer insight into potential collateral effects on bacterial fitness and prove useful for developing novel treatment strategies based on evolutionary principles.

In this study, we investigated the evolutionary dynamics of resident gut bacteria during intestinal inflammation using an established gnotobiotic mouse model of IBD ^26^. Through the monocolonization of both *Interleukin 10*-knockout and wildtype mice with a single *E. coli* strain (NC101), this setup allowed us to track genetic and phenotypic changes over the course of intestinal inflammation and to evaluate their potential clinical relevance. By employing multiomic analysis and high-throughput phenotypic screening, we identify genetic changes in bacteria, alterations in the metabolome, and differences in numerous phenotypic traits among bacterial populations that are specifically associated with the evolution in inflamed *Il10*-knockout mice. Remarkably, among the phenotypic changes observed in inflammation-adapted bacteria are sensitivities to antibiotics with a known therapeutic value in IBD. These results further confirm the importance of understanding bacterial adaptation to inflammation and suggest its more widespread study of patients as a means to develop novel treatment approaches.

## Results

### Gnotobiotic model of intestinal inflammation

In order to capture the evolutionary dynamics of bacteria evolving in the context of intestinal inflammation, we implemented a previously established gnotobiotic model, for which inflammation develops upon colonization with the *E. coli* NC101 strain in IL10-deficient (*Il10^−/−^*; herein “KO”), but not wild type (WT) mice ^26^. We performed two independent *in vivo* experiments (see Methods), in which WT (total N=14) and KO (total N=11) mice were monocolonized and monitored over a period of 12 weeks, with longitudinal sampling of feces for downstream multiomic analyses (Fig 1A; Table S1, see Methods). Inflammation was monitored via levels of lipocalin-2 in feces, which steadily and significantly increased in KO mice, but not in WT mice, already at one week post-inoculation (Fig 1B, Tables S2 and S3, Wilcoxon signed-rank test Benjamini-Hochberg-corrected *P* < 0.05). Histopathological assessment of colonic tissue at the endpoint reveals significantly higher pathology scores in KO mice than in WT mice (Fig 1C, Kruskal-Wallis H test Benjamini-Hochberg-corrected *P* < 0.05), further confirming that inflammation was specific to KO mice. The bacterial load in the fecal samples was stable throughout the experiment, as measured by CFU counts normalized by wet feces weight (Fig S1, Tables S4 and S5).

**Figure 1.**
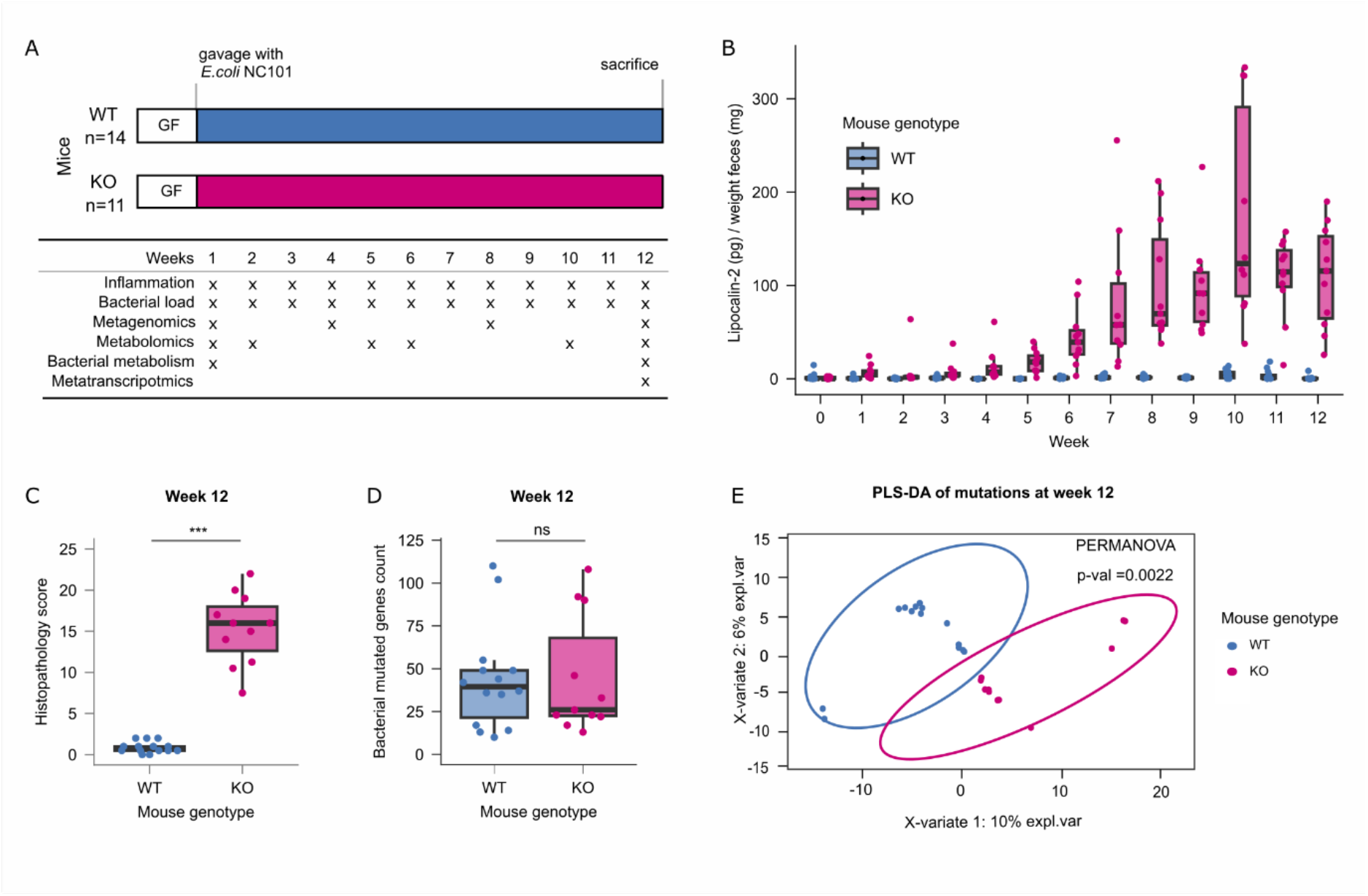
Application of a mouse model of IBD to test the effect of gut inflammation on the evolution of NC 101. **A** Schematic of the experimental setup. Germ-free WT and *Il10*^-/-^ mice were monocolonized with *E. coli* NC101. Fecal samples were collected from each mouse at the indicated time points and analyzed as described. **B** Fecal lipocalin-2 levels from the WT and *Il10*-/- mice. Lipocalin-2 was measured using the Mouse Lipocalin-2/NGAL DuoSet ELISA and normalized to feces weight. Each dot represents one mouse at each sampling point. Results of Wilcoxon signed-rank test of lipocalin-2 concentrations at each time point are reported in Table S3. Differences were considered statistically significant at Benjamini-Hochberg-corrected *P* <0.05. **C** Boxplots of the histopathology scores of the colon tissue of mice after sacrifice at the end of the experiment. Each dot represents a single mouse. Differences were considered statistically significant using the Kruskal-Wallis H test at Benjamini-Hochberg-corrected *P* <0.05. **D** Number of *de-novo* mutations in the bacterial populations at week 12 in WT and KO mice compared with the reference genome. Each dot represents a bacterial population from a single mouse. Differences were considered statistically significant using the Kruskal-Wallis H test at Benjamini-Hochberg-corrected *P* <0.05. **E** Partial least squares-discriminant analysis of the *de-novo* mutated genes in the evolved populations of *E. coli* NC101 at week 12 in WT and KO mice compared to the reference genome. Differences were considered statistically significant at a PERMANOVA *P* < 0.05.

### Bacterial populations show genetic diversification in healthy and inflamed mice

To assess the genetic diversification of the bacterial populations during the evolution experiment, we performed shotgun sequencing on each single inoculum and all bacterial populations at weeks 1, 4, 8, and 12 and identified *de-novo* mutations (i.e., mutations that were not present in an inoculum; see Methods) by comparison with the genome of the ancestral strain. No statistically significant difference was observed in the number of mutations in the bacteria from KO and WT mice at any point in the experiment, suggesting that differences in the intestinal environment in which the populations evolved did not exert a measurable effect on the number of mutations (Fig S2, Table S6). One week after gavage, the bacterial populations showed a median number of 87 *de-novo* mutations, with a subsequent reduction at the following timepoints, although the differences are not significant (median at week 4=34, at week 8=45 and at week 12=36, Fig S2, Tab S7 Wilcoxon signed rank test, corrected *P* > 0.05). Comparison of the evolved populations at week 12 with the ancestral strain reveals a median number of 39 mutations per mouse in WT and 26 in KO mice (Fig 1D). This indicates that the evolved bacterial populations in both groups of mice genetically differ from the ancestral strain. Although the unphased shotgun data do not enable us to directly analyze the role of genetic hitchhiking, Table S8 indicates the number of mutations at week 12 already present at each earlier time point. Interestingly, there remains a substantial (on average approx. 60% in each mouse genotype) proportion of *de-novo* mutations that arose between week 8 and 12, which suggests that a large proportion of the mutations cannot be explained by hitchhiking from standing genetic variation at earlier time points.

To test for overall differences in the *de-novo* mutations that accumulated in the bacterial populations that evolved in the WT and KO mice, we performed a partial least-squares discriminant analysis (PLS-DA) on all mutated loci (genes or intergenic regions differing in their nucleotide sequence compared to the inocula). We find a clear and significant distinction between the populations that evolved in the two mouse genotypes (Fig 1E, PERMANOVA *P* = 0.0022). The same result was observed when comparing single *de-novo* mutated genomic positions between evolved bacterial populations from WT and KO mice (Fig S3, PERMANOVA *P* = 0.0016).

### Single nucleotide polymorphism differentiates evolved bacteria of all inflamed mice from evolved bacteria of healthy mice

To identify candidates for genetic changes that may be specifically selected in an inflamed environment, we focused on parallel mutations (i.e., genes or intergenic regions mutated in at least two mice that were gavaged with an independent inoculum; see Methods) and their associated functions. Analysis of the KEGG pathways encompassed by these loci reveals seven pathways unique to the bacterial populations from inflamed mice (Fig 2A).

**Figure 2.**
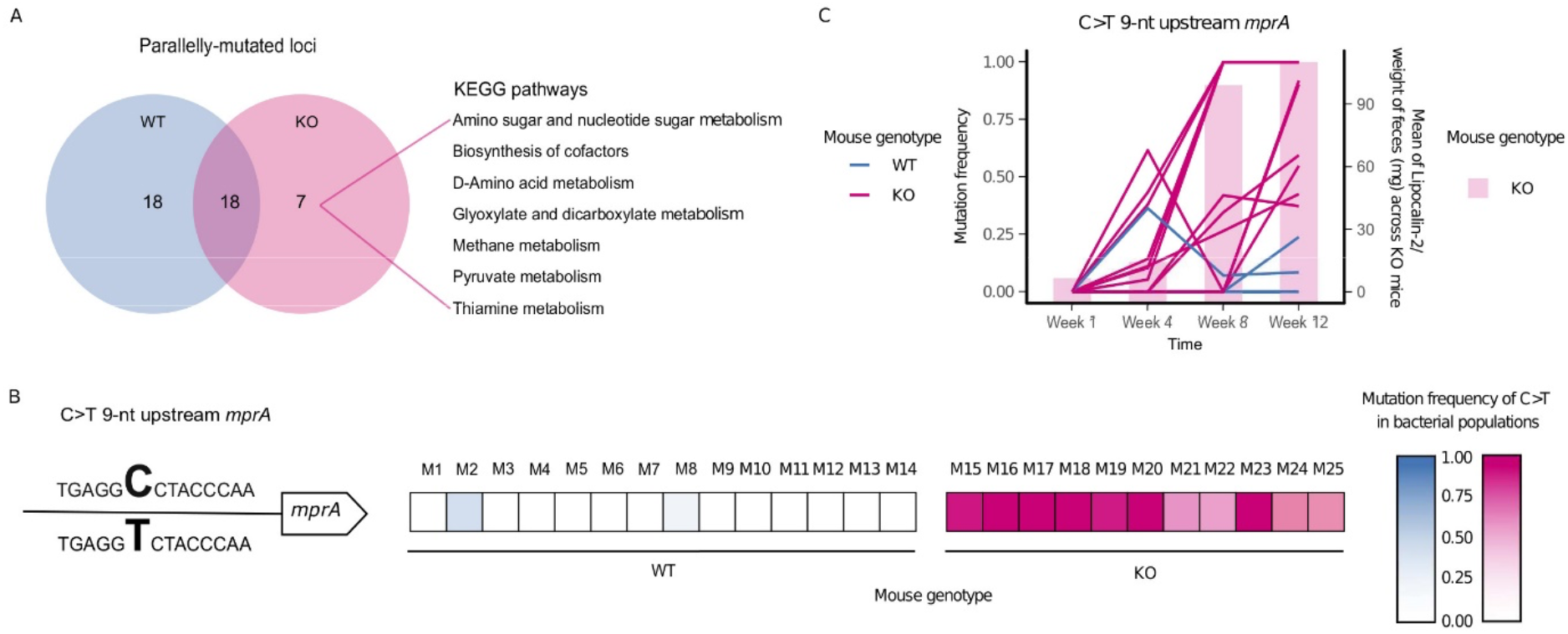
Parallel evolution of *E. coli* in inflamed mice revealed a disease-specific genetic signature. **A** Parallelly mutated genes were classified based on KEGG pathways; **B** An intergenic C>T mutation 10-nt upstream of *mprA* was the unique parallel mutation significantly more abundant in populations evolved in the inflamed gut (Wilcoxon signed-rank test Benjamini-Hocheberg corrected *P* < 0.005). The heatmap shows the frequency of the mutation in bacterial populations from each mouse. **C** Mutation frequency (left axis) of the intergenic C>T mutation 10-nt upstream of *mprA* and mean of lipocalin-2 concentrations across KO mice at different timepoints during the *in-vivo* evolution experiment. Each line represents a population from a single mouse. Each bar represents the mean value of the normalized concentration of lipocalin-2 in fecal samples from KO mice.

These pathways include those involved in amino acid metabolism, pyruvate metabolismall of which have been associated with IBD in clinical settings ^27,28^. In addition to the pathways affected only in bacterial populations from the inflamed mice, 18 pathways are affected only in populations from healthy mice, and 18 other pathways are mutated in populations from both healthy and inflamed mice (Fig 2A).

Univariate analysis reveals that only one particular locus is mutated at significantly higher frequencies in evolved bacterial populations from inflamed than healthy mice (Wilcoxon signed-rank test, corrected *P* = 0.001, Table S9), which is a single nucleotide polymorphism (SNP) (C>T) nine nucleotides upstream of the gene *mprA* (position 3009211) (Fig 2B). It is found in 11/11 KO mice at the endpoint with a frequency between 0.373 and 1, compared to 2/14 WT mice with a frequency between 0.085 and 0.239. Two other mutations are detected in the same intergenic region, although in a few populations only. The bacteria from one WT mouse harbors a C>A mutation at the same position (3009211) with a frequency of 0.052, whereas those from one inflamed mouse harbors a G>T SNP at a nearby position (3009194) with a frequency of 0.611.

To further investigate the most frequent C>T mutation, we analyzed its frequency over the course of the experiment (weeks 1, 4, 8, and 12). At week 1 the mutation is not present or below the detection threshold (minimum frequency of 0.05; see Methods), while at week 4, it is observed in 7/11 KO mice with a maximum frequency of 0.616, in comparison to 1/14 WT mice with a frequency of 0.364 (Fig 2C). After a steady increase in frequency at weeks 8 and 12, the mutation reaches fixation or near fixation in 7/11 KO mice and is also present at intermediate frequencies in the remaining 4/11 KO mice (Fig 2C). Importantly, the increase in frequency of the C>T mutation strongly coincides with the increase in inflammation as measured by lipocalin-2 levels, both in KO mice alone and when including all mice (Fig 2C, Fig S4; Spearman’s rank correlation rho *P* = 0.00005 for KO mice alone and *P* = 3.005e-11 for all mice).

Interestingly, closer inspection of the intergenic region between the hypothetical protein_02879 and the gene *mprA* reveals that this genomic region is a mutational hotspot, with six others positions in this region having mutated over the course of the *in vivo* experiment (Fig S5).

Taken together, we identify a C>T change at position 3009211 in the *E. coli* genome, which is a parallel mutation significantly associated with intestinal inflammation. This mutation is located upstream of *mprA* (also known as *emrR*), a transcriptional repressor of several genes, including those encoding the efflux pump EmrAB and AcrAB, and a putative outer membrane porin OmpC (also known as NmpC)^29–31^.

### Bacterial populations show significantly different expression profiles in healthy and inflamed mice

To quantify the expression of genes that are part of the *mprA* regulon (*mprA*, *emrAB*, *acrAB*, and *ompC*)^29–31^, as well as gain an overall view on differences in gene expression in WT-compared to KO-evolved bacterial populations, we perform metatranscriptomic sequencing of all week 12 bacterial populations. To test for overall differences in the gene expression, we perform a partial least-squares discriminant analysis (PLS-DA) on all genes, using RPKM (Reads per Kilobase per million Mapped Reads) as a proxy for gene expression. We find a clear and significant difference in overall gene expression between populations according to mouse genotype (Fig S6A, PERMANOVA P < 0.00005). Differential expression analysis performed with Deseq2 reveals 2,219 genes that are up- or down- regulated in one of the two conditions (1,022 upregulated for the populations from the healthy gut and 1,197 for the population from the inflamed gut; Benjamini-Hochberg corrected *P* <0.05, Tab S10). Analysis of the KEGG pathways encompassed by these upregulated genes reveals 25 pathways unique to the bacterial populations from inflamed mice (Fig S6B). These pathways include Ribosome, Aminoacyl-tRNA biosynthesis, Glutathione metabolism, RNA degradation, Folate biosynthesis, Sulfur relay system, Arginine biosynthesis, Protein export, Pantothenate and CoA biosynthesis, and Homologous recombination, among others (Tab S11). Interestingly, *mprA* and *ompC,* which can be putatively regulated by *mprA*^30^, are significantly more expressed in the populations that evolved in the inflamed gut (Tab S10; Fig S6C).

### Increase in fitness of evolved bacteria in an inflammatory environment

As parallel mutations are a strong indication of adaptation by natural selection, we hypothesized that the evolved bacterial populations may have a fitness advantage in the environment in which they evolved, and thus next focused on the phenotypes present in the evolved bacterial populations. To test whether evolution in the inflamed mouse gut confers a fitness advantage to *E. coli*, we performed “reciprocal transplant” experiments based on *ex vivo* assays using filtered cecal content from inflamed versus non-inflamed mice, collected after sacrifice (referred to subsequently as “*ex vivo* media”). Populations of *E. coli* from fecal samples collected at week 12 from a subset of mice (N = 5 KO mice and N = 5 WT mice) were cultured in *ex vivo* media derived from both inflamed and non-inflamed mice. While no significant difference is observed between bacteria from WT and KO mice when grown in non-inflamed *ex vivo* media (Kruskal-Wallis test *P* = 0.9168; Fig 3A), bacteria from KO mice grow significantly better than those from WT mice in the inflamed *ex vivo* media (Kruskal-Wallis test *P* = 0.009023). Notably, bacteria from KO mice grow significantly better in the inflamed *ex vivo* media than in the non-inflamed *ex vivo* media (Kruskal-Wallis test *P* = 0.0472), while bacteria from WT mice show no difference in growth in the two media (Kruskal-Wallis test *P* = 0.4647). Thus, bacteria that evolved in the inflamed gut possess a fitness advantage in the inflamed gut environment, suggesting that evolution in the inflamed gut results in specific adaptation to that environment.

**Figure 3.**
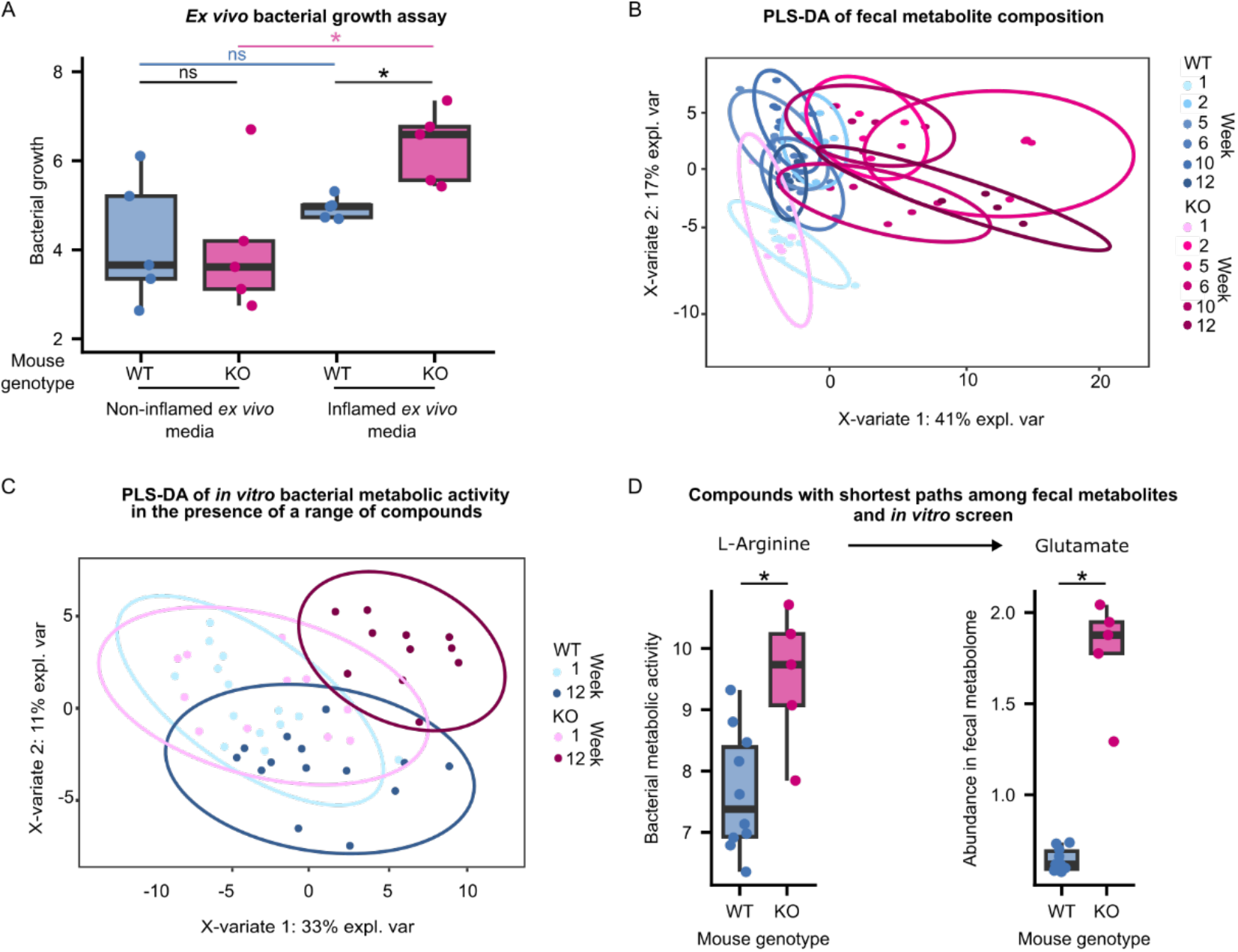
Phenotypic characterization of evolved *E. coli* and the inflammatory environment in which they were selected. **A** Evolved populations from a subset of mice (n = 5 KO mice and n = 5 WT mice) were cultured in media derived from the cecal content collected from healthy (WT) and inflamed (KO) mice after sacrifice. The area under the curve (AUC) calculated from growth curves measured during the growth of evolved populations from each mouse in inflamed and healthy cecal contents is shown. Each dot represents the mean of three independent replicates. * *P* < 0.05 (Kruskal-Wallis H test); ns, not significant. **B** Partial least squares discriminant analysis (PLS-DA) of the fecal metabolite abundances at different time points during the evolution experiment. Differences were considered statistically significant at PERMANOVA *P* < 0.05 **C** Bacterial metabolism was measured *in-vitro* in a range of compounds. Measurements were performed on bacterial populations derived from fecal samples collected at weeks 1 and 12 of the *in-vivo* evolution experiments. Partial least squares discriminant analysis (PLS-DA) was performed on the area under the curves (AUCs) of the measured metabolic activity of the evolved bacterial populations from all mice in the range of compounds. Differences were considered statistically significant at PERMANOVA *P* < 0.05 **D** Compounds from the fecal metabolome and the *in vitro* screen with the shortest path were inspected. One of the pairs with the shortest path was glutamate-arginine. Glutamate was enriched in the fecal metabolome of KO mice, whereas KO-adapted bacteria showed higher metabolic activity in the presence of arginine. Each dot represents the abundance of the compound in the fecal sample from a mouse (for glutamate) or the mean metabolic activity of the bacterial population from a mouse in the presence of L-arginine from two independent measurements (for arginine). * *P* < 0.05, (Kruskal-Wallis test at Benjamini-Hochberg-corrected), ns, not significant.

### Inflamed gut environment displays an altered metabolomic profile

To gain a better understanding of the environment to which these bacteria are adapted, we performed metabolomic analysis of fecal samples spanning the experimental time course (weeks 1, 2, 5, 6, 10, and 12) using 1H-NMR in mice from one of the two experiments. While WT and KO mice initially display very similar fecal metabolomes (PERMANOVA *P* = 0.754), with time, the fecal metabolic composition begins to vary concurrently with the onset of inflammation and the frequency of the C>T SNP upstream of *mprA* (Fig 3B, 1B, 2C). By week 12 of the evolution experiment, WT and KO mice exhibit notable differences in their fecal metabolomes, as revealed by PLS-DA (PERMANOVA *P* = 0.001; Fig 3B, Fig S7). Among the 77 metabolite features initially found in the samples from week 1, none show significantly different abundances between the WT and KO mice (Table S12). In contrast, in the week 12 samples, 59 of the 77 detected features exhibit significant differences in abundance between WT and KO mice (Benjamini-Hochberg corrected *P* < 0.05; Table S13). Notably, 42 are significantly more abundant in KO mice than in WT mice, including all detected amino acids that were significantly enriched in KO mice based on KEGG pathway analysis of parallelly mutated loci. Among the 17 enriched features in the WT mice, most are putatively annotated as (poly-)saccharides (Fig S8). Notably, variations in fecal metabolic composition at week 12 could arise from changes in both the host, linked to inflammation-related changes, and the bacterial population, associated with bacterial adaptation to inflammation.

### Phenotypic profiling of evolved bacteria compared to ancestor

Having established that the environment in which the bacteria evolved significantly differs according to the mouse genotype, we next focused on specific bacterial traits that may have changed as a consequence of adaptation. Accordingly, we tested the metabolic activity of the *E. coli* populations using the Biolog GENIII test panel, which comprises 94 unique biochemical tests, including a range of compounds (e.g., sugars, amino acids, and short-chain fatty acids), conditions (e.g., different pH and salt concentrations), and chemical sensitivities (e.g., antibiotics; see Methods). Similar to the pattern observed for the fecal metabolome, the overall bacterial metabolic activity across all tested conditions is similar among populations derived from the WT and KO mice at week 1 of the experiment (PERMANOVA, *P* = 0.519; Fig S9), as also evidenced by the overlapping of the groups in the PLS-DA based on the overall metabolic activity observed in the BIOLOG GENIII tests (Fig 3C). However, by week 12, the overall pattern of metabolic activity across all tested conditions significantly differs between the bacteria from KO and WT mice (PERMANOVA, *P* = 0.005, Fig S9). These results suggest that adaptation to the inflamed intestine results in significantly altered bacterial metabolism and the ability to grow in the presence of different inhibitors. Among the compounds included in this screen, bacteria from WT and KO mice from week 1 showed significantly different metabolic activity in only one compound (Table S14). In contrast, bacteria from WT and KO mice at week 12 display significantly different metabolic activities in the presence of 31 compounds (Wilcoxon signed-rank test, corrected *P* < 0.05; Table S15, Fig S10).

To determine whether the differences in growth observed among the 31 compounds at week 12 may be related to the altered metabolomic environment of inflamed mice, we compared our BIOLOG GENIII results to the compounds found to be enriched in the fecal metabolome of the KO mice. Interestingly, metabolomics analysis revealed a significant enrichment of the amino acid histidine in the feces of KO mice compared to that of WT mice (Fig S8). In our *in vitro* metabolic assay, we found that bacteria adapted to the inflamed intestine had a significantly higher ability to metabolize histidine than bacteria adapted to the healthy intestine (Fig S10). Thus, adaptation to an environment enriched in histidine may have conferred a higher metabolic activity in the presence of this amino acid.

To further explore how the results of our *in vitro* phenotypic analyses may relate to the fecal metabolomic profile observed *in vivo*, we performed a shortest path analysis on the metabolic network of the genome-scale metabolic model of *E. coli* NC101 ^32^ using metabolomic data and data from the *in vitro* metabolic activity screen (BIOLOG plates; see Methods). Pairs of compounds from the two datasets that are converted easily to each other by means of pathway length include glutamate and L-arginine (Fig 3D, Table S16). Glutamate is significantly more abundant in the feces of the inflamed mice (Fig S8), whereas bacteria adapted to the inflamed intestine are capable of significantly higher metabolism in the presence of L-arginine (Fig S10). This result suggests that the bacteria may be converting arginine to glutamate, resulting in the enrichment of glutamate in KO mice (Fig 3D, Table S16). A second pair of compounds showing the same pattern are taurine and L-arginine. However, analysis of the potential pathways between these compounds through flux variability analysis revealed that taurine cannot be produced by *E. coli* NC101 under any of the tested conditions (Fig S11). Additionally, taurine abundance increased steadily over time only in KO mice (Fig S12). Thus, taurine is most likely to be produced by the host.

Finally, we examined 31 significant differences in metabolic activity observed between bacteria from WT and KO mice at week 12, in light of their potential clinical value. A first compound of interest is N-acetyl beta-D mannosamine (NADM), which is a precursor to sialic acid and has been shown to promote *E. coli* colonization of inflamed gut ^33^. Our *E. coli* populations show higher levels of metabolic activity in the presence of NADM after adaptation to the inflamed mouse gut. In contrast, *E. coli* evolved in the non-inflamed mouse gut show no such change (Fig 4). These findings suggest that increased virulence may occur in the context of adaptation to an inflamed environment.

**Figure 4.**
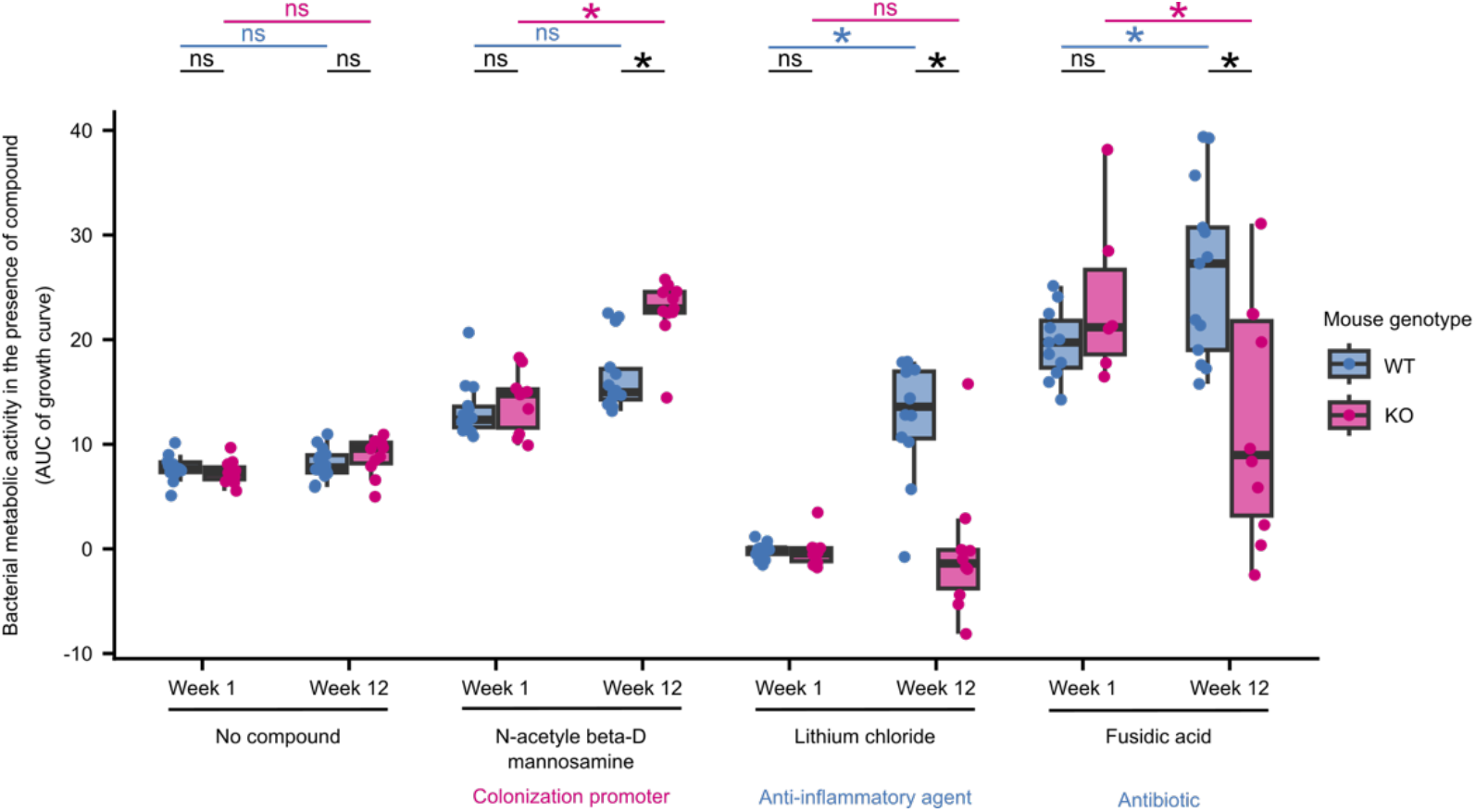
*E. coli* adapted to the inflamed gut show phenotypes with clinical relevance. Metabolic activity of bacteria isolated at weeks 1 and 12 of the evolution experiment in the presence of three selected compounds. Area under the curve was calculated from growth curves measured during the growth of evolved populations in the presence of each compound. Each dot represents the mean of two independent replicates of the metabolic activities of the *E. coli* population from a mouse measured in two independent experiments. *, *P* < 0.05 (Kruskal-Wallis test at Benjamini-Hochberg-corrected). The clinical relevance of the three compounds is indicated.

Another interesting candidate is lithium chloride, which is known to have anti-inflammatory effects via the inhibition of a key host regulator of inflammation, glycogen synthase kinase-3 beta ^34^. In our experiment, bacterial populations evolved in WT mice display improved metabolic capacity in the presence of lithium chloride compared to the ancestor, while no such change is observed among those evolved in the KO mice. Thus, our results suggest that an inflamed intestinal environment may prevent the acquisition of this phenotype in *E. coli*.

A final promising category of candidates are antibiotics, including fusidic acid, vancomycin, and lincomycin. When considering the loading plot of the PLS-DA of our Biolog GENIII data (Fig S13; reporting which compounds contribute the most to the separation between groups), all three of these antibiotics contributed to the overall significant difference in the metabolic activity of the bacteria from WT and KO mice (Fig 3C, Fig S13). Furthermore, fusidic acid is also significantly different between bacteria from the two mouse genotypes in the univariate analysis (Wilcoxon signed-rank test, corrected *P* < 0.05). Fusidic acid is an antibiotic that targets gram-positive bacteria, and is commonly used to treat skin infections ^35^. Interestingly, it was also shown to be effective in reducing disease activity in a small number of patients with Crohn’s disease, which is thought to be due to its immunosuppressive properties ^36,37^. We observe *E. coli* to display decreased metabolic activity in the presence of fusidic acid after adaptation to the inflamed intestine but increased metabolic activity after adaptation to the healthy intestine. These results suggest a potential trade-off between adaptation to inflammation and resistance to fusidic acid, where *E. coli* adapted to inflammation also show lower resistance to fusidic acid.

In summary, we observe widespread phenotypic differences among the bacterial populations that evolved in the inflamed intestines of KO mice compared to non-inflamed WT mice. These results relate to differences in the metabolome between these two intestinal environments and include phenotypes that carry the potential for exploitation in a clinical setting.

## Discussion

In this study, we describe the findings of an *in-vivo* bacterial evolution experiment using a gnotobiotic mouse model of IBD. We find that *E. coli* evolved in the inflamed mouse gut accumulated specific genetic changes and that these changes confer a fitness advantage in the inflamed intestinal environment, which significantly differs in its metabolome. Furthermore, we show that *E.coli* populations in the gut of healthy and inflamed mice have a distinct metatranscriptomic profile. Finally, we showed that adaptation to the inflamed intestine resulted in several phenotypic differences that may be clinically relevant, such as differential tolerance to antibiotics.

Recent studies have highlighted how strain-level changes in members of the gut microbiome can play a crucial role in the adaptation of the gut microbiome to novel conditions ^14,38,39^. However, the effects of inflammation on the evolution of gut commensals remain largely unexplored. Barreto et al. (2020) followed the adaptation of *E. coli* in mice of different ages *in vivo* ^14^. They showed that the aged mouse gut, which also showed high levels of inflammation, was a more stressful environment for *E. coli*, resulting in a higher number of mutations and more severe selective pressure on commensals, particularly in bacterial loci associated with stress-related functions. Notably, we find no difference in the number of mutations acquired by *E. coli* in inflamed mice, suggesting that other factors associated with aging may contribute to the increased bacterial mutation rate observed by Barreto et al. (2020) ^14^.

More recently, Tawk et al. (2023) conducted a study of mice monocolonized with *Bacteroides thetaiotaomicron* that were subsequently infected with *Citrobacter rodentium* and developed inflammation. In this setting, a single-nucleotide variant of *B. thetaiotaomicron*, which had a higher tolerance to oxidative stress than the ancestral variant, underwent selective sweeps and dominated the intestinal community ^39^. Although the experimental setup and duration of inflammation differ between this study and the current study, our candidate mutation is also known to be associated with oxidative stress ^40^. This is consistent with the observations of Barreto et al. (2020) and Tawk et al. (2023) ^14,39^, suggesting that oxidative stress is a key selective pressure in an inflamed environment.

Moreover, oxidative stress plays a major role in the pathophysiology of IBD ^41^. Interestingly, this is also confirmed by our metatranscriptomic analysis. In particular, 14 of the genes upregulated in KO mice belong to the pathway of glutathione metabolism (eco00480), and previous studies demonstrated that glutathione metabolism is involved in protection against oxidative stress ^42,43^. In addition, Sakamoto et al. reported *mprA* to be involved in resistance to oxidative stress, together with the activity of *gshA* (glutathione synthase), which is also upregulated in our populations^40^. Additionally, our metatranscriptomic data confirm the findings of Tawk et al., suggesting a role for vitamin B6 metabolism ^39^. We observe not only the upregulation of nine genes belonging to the vitamin B6 metabolism (eco00750), but also a significantly higher concentration of aspartate, a substrate of vitamin B6-dependent enzymes, in the feces of the inflamed mice^39^.

Importantly, many of our findings are consistent with the observations made in clinical IBD settings. First, the KEGG pathways affected by parallel mutations specific to the inflamed mice (Fig 2A) include D-amino acid-, pyruvate-, and thiamine metabolism. This is in line with previous reports that the basic metabolism is reduced in IBD ^27,44^.

Second, the metabolomic profiling of the fecal samples reveals that KO mice have significantly higher levels of many amino acids, which was also reported in IBD patients ^45^. Furthermore, the results of our shortest path analysis suggest that bacterial production of glutamate from arginine may underlie the enrichment of glutamate in the inflamed gut. Although taurine-arginine was also one of the shortest paths we found, taurine could not be produced by *E. coli* according to the flux variability analysis. Thus, taurine is most likely to be produced by the host. Taurine is known to be a promoter of colonization resistance, and infection has been shown to prime the microbiota against subsequent infections by inducing host production of taurine ^46^. Indeed, taurine was also reported to ameliorate inflammation in rat models of inflammation ^47^. Furthermore, taurine is a substrate for the microbiota-driven production of hydrogen sulfide ^48^, and arginine is a precursor to polyamines that can protect against reactive species such as hydrogen sulfide ^49^. Thus, bacterial adaptation for improved metabolic activity in the presence of arginine may be a response to the host-induced inflammatory changes in the intestine.

Third, *E. coli* adapted to the inflamed mouse gut also showed some clinically relevant phenotypes (Fig 4). N-acetyl beta-D mannosamine (NADM) is a precursor of polysialic acid, a pathogenic determinant, and studies have shown that it can promote colonization of the inflamed intestine by pathogenic *E. coli* ^33,50^. As NADM is a crucial player in mediating pathogenic activity in the inflamed intestine, the improved metabolic activity of inflammation-adapted *E. coli* observed in the presence of NADM may be a phenotype associated with virulence. This is consistent with reports of virulence-associated phenotypes, such as hypermotility, selected during the evolution of an adherent-invasive *E. coli* in the inflamed mouse gut ^13^. Furthermore, lithium chloride is known to inhibit the activity of glycogen synthase kinase 3-β, a master regulator of host chronic intestinal inflammation mediated by toll-like receptors, and has been used as a treatment in a mouse model of IBD ^34,51^. However, the effect of lithium on the microbiome has not yet been elucidated. *E. coli* tolerance to lithium ions is regulated through antiporters, and proline has been shown to induce the uptake of lithium ions by *E. coli* ^52,53^. Our finding that adaptation to the healthy mouse intestine confers improved metabolic capacity in the presence of lithium chloride, while adaptation to the inflamed intestine does not (Fig 4), implies that lithium may affect both the host and microbiome. This may also imply that adaptation to the healthy intestine affects the antiporter-dependent detoxification system of *E. coli*, resulting in improved ion tolerance.

Lastly, we found that bacterial adaptation to the inflamed gut decreases metabolic activity in the presence of fusidic acid, while adaptation to the healthy mouse gut improved metabolic activity in the presence of fusidic acid (Fig 4). Fusidic acid is an antibiotic with T cell-specific immunosuppressive effects that also stimulate gastric mucus secretion. Furthermore, it was successfully applied to alleviate inflammation in rats, as well as to treat selected patients with Crohn’s disease for whom conventional treatment was ineffective ^36,37^. However, the effect of fusidic acid on the gut microbiome in IBD has not yet been studied. Our results imply that bacterial adaptation to the inflamed gut can result in a trade-off for resistance to fusidic acid. Importantly, such trade-offs are a fundamental concept in evolutionary medicine, where increased fitness in one context results in a consequent decrease in fitness in another context, and are proposed as possible potent therapeutic strategies against antibiotic resistance and cancer ^18,54^. Thus, the therapeutic effect of fusidic acid observed in patients with CD is due to trade-offs in microbiome adaptation to the inflamed gut and/or its immunosuppressive effects on the host. Interestingly, it is known that lipocalin-2 plays a role in iron-sequestration, which has been reported to have a role in antibiotic resistance and sensitivity^55^. It is thus worth noting that there is no enrichment of genes involved in iron metabolism among the mutated genes in the populations evolved in the inflamed gut, as well as among the upregulated genes in these populations.

Current treatment options for IBD are largely limited to the treatment of inflammation with a chronic risk of relapse. Corticosteroids, aminosalicylates, and immunosuppressive agents are the conventional drugs of choice, but the safety and efficacy of novel emerging strategies remain unclear (reviewed in ^56^). Given that disease-specific aspects of the microbiome have been shown to remain stable over long periods in IBD patients despite treatment ^57^, a persistent microbial influence on the intestinal environment may be a key risk factor for relapse. Thus, our results highlight new avenues of research involving evolution-informed therapeutic strategies that exploit trade-offs to either prevent adaptation to inflammation and/or help restore desirable ancestral traits in the microbiome.

While our results provide valuable insights into the potential role of evolution-informed therapeutic strategies, it is important to acknowledge that it remains unknown whether the same or similar genotypic and phenotypic changes would be observed in the context of a complex microbial community as present in the human gut microbiome. Future work should therefore include similar *in vivo* evolution experiments using a complex or synthetic microbial community.

## Methods

### Bacterial strains

*E. coli* NC101 strain was obtained from Balfour Sartor, University of North Carolina, Chapel Hill, NC, USA ^58^. *Escherichia coli* NC101 is a mouse strain isolated from the intestine of a WT 129S6/SvEv mouse raised under specific-pathogen-free (SPF) conditions^58^. The strain was cultured at 37 °C in Luria Bertani (LB) medium with continuous shaking. The ancestor strain for the *in vivo* experiment was obtained by plating the overnight culture on LB agar and picking single colonies to be used as inocula for the *in vivo* evolution experiment.

### Mouse model

Two independent experiments were conducted to study *E. coli* adaptation to the inflamed gut (Table S1). In the first experiment, “Exp1”, a single inoculum was used for all mice. In order to ensure that mutations present in a common inoculum could not be falsely identified as “parallel” mutations, the second experiment, “Exp2” was performed with independent inocula for each mouse. Accordingly, our definition of a parallel mutated gene requires it to be mutated in at least two mice that were gavaged with a different inoculum. Germ-free (GF) C57BL/6NTac (WT) and C57BL/6NTac-Il10em8Tac (KO) male mice were purchased from Taconic Biosciences (Silkeborg, Denmark) and housed in the Germ-Free Animal Facility at the Max Planck Institute for Evolutionary Biology (Ploen, Germany). GF mice were maintained in sterile isolators (MB-10, Quip Laboratories, Delaware, USA) and fed sterilized 50 kGy V1124-927 Sniff (Soest, Deutschland). The animals were allocated to independent cages with a maximum of four mice per isolator until they reached an age of 12 weeks. Detailed information regarding the mice used in this study is presented in Table S1. Initially, 19 WT and 15 KO mice were mono-colonized with the *E. coli* NC101 strain. Only mice that survived the duration of the experiment (12 weeks; N = 14 WT mice and N = 11 KO mice) were included in the study. The study was performed in accordance with the approved animal protocols and institutional guidelines of the Max Planck Institute for Evolutionary Biology, Plön. Mice were maintained, and experiments were performed in accordance with FELASA guidelines and German animal welfare law (Tierschutzgesetz § 11; permits from Veterinäramt Kreis Plön: 1401-144/PLÖ-004697 and Veterinäramt Kreis Kiel: 244-509017/2018(107-11/18)).

The strains used as inocula were diluted in sterile PBS, and 200 μl (equivalent to 1×10^8 bacteria) were gavaged using a sterile gavage needle (Reusable Feeding Needles 18G, Fine Science Tools, Heidelberg, Deutschland). Fecal pellets were collected in a sterile manner twice a week to study the evolution of *E. coli* for a total of 24 samples collected per mouse and once a week for metabolomic investigation. An overview of the sampling plan is presented in Fig 1A.

### Histopathological evaluation

Mice were sacrificed at week 12, and colon tissue was collected and arranged to form a Swiss-roll ^59^. Hematoxylin and eosin-stained sections of colonic tissue Swiss rolls were scored by two independent researchers in a blinded manner using the scoring system described by Adolph et al. (2013) ^60^. The score is composed of five sub-scores: mononuclear cell infiltrate, crypt hyperplasia, epithelial injury or erosion, polymorphonuclear cell infiltrates, and transmural inflammation. Each of the first four sub scores was awarded a score from 0 to 3, whereas transmural inflammation was scored from 0 to 4, with a higher score indicating a more severe level of inflammatory activity. The sum of the sub scores was then multiplied by a factor based on the percentage of affected bowel length (1= < 10%; 2= 10-25%; 3= 25-50%; 4= >50%).

### Feces processing

Feces weight and consistency were recorded. Feces were homogenized in 1.5 ml sterile PBS on a horizontal vortexer (Vortex Mixer Modell Vortex-Genie® 2, Scientific Industries, Bohemia, NY, USA) at maximum speed for 30 min, then separated in three aliquots of 500 μl. Each aliquot was centrifuged at 10,000 rpm for 5 min. The supernatants were transferred to a new tube and stored at −20 °C for subsequent lipocalin-2 concentration measurement. One pellet aliquot was resuspended in 500 μl of PBS and used to prepare a 12-point 1/10 dilution series in a 96-well plate. Ten μl were plated onto an LB agar plate. Plates were incubated overnight at 37°C and colonies were counted to estimate the bacterial load. The bacterial load in the fecal samples was calculated by normalizing CFU counts by wet feces weight. The second pellet aliquot was resuspended in 1 ml RNAlater and stored at +4 °C for 24 h, after which the tube was centrifuged at 10,000 rpm for 5 min to remove the RNAlater, and the pellet was stored at −20 °C for subsequent nucleic acid extraction and sequencing. The final pellet aliquot was resuspended in 500 μl of LB containing 20% glycerol and stored at −70 °C for further phenotypic investigation.

### Lipocalin-2 quantification

Lipocalin-2 concentration in the supernatants was measured using the commercial kit Mouse Lipocalin-2/NGAL DuoSet ELISA (R&D Systems, Minneapolis, MN, USA) for mouse Lipocalin-2 (DY1857). Testing was performed according to the manufacturer’s instructions. The samples were diluted 1:10 and added to the plate. The optical density of each well was determined using a plate reader (SPARK, Tecan, Tecan, Männedorf, Switzerland) at 450 and 540 nm. Lipocalin-2 concentrations were normalized to the weight of feces for each sample.

### Shotgun sequencing

Total DNA and RNA were extracted using the ZymoBIOMICS DNA/RNA Mini Kit (Zymo Research, Freiburg, Germany), following the manufacturer’s instructions from fecal pellets collected at weeks 1, 4, 8, and 12, as well as from the *E. coli* NC101 cultures that served as inocula. Shotgun metagenomic sequencing was performed on four Illumina Nextseq (HighOutput 300 cycles) sequencing runs at the Max Planck Institute for Evolutionary Biology (Plön, Germany).

Raw reads were filtered and trimmed to ensure good quality using Cutadapt (version 3.2, ^61^). First, any pair containing Ns, homopolymers (10 nucleotides or more), or those longer than 151 bp were discarded. Sequences were trimmed with a quality cutoff of 25 at both ends for both reads, Illumina adapters were removed, and sequences shorter than 50 bp were discarded. Good quality sequences were then filtered to exclude mouse sequences (mouse_C57BL) using KneadData ^62^. A final filter was applied to remove any adapter leftovers using Trimmomatic ^63^ and sequences shorter than 105 bp were discarded. Mutations were identified using Breseq ^64^ by comparing the obtained metagenomes with the reference genome of *E.coli* NC101 (PRJNA596436) using default options (i.e. mutations identified by a threshold frequency > 0.05) and by subtracting the mutations that were detected in the respective inoculum. Gdtools was used to compare mutations in the samples.

### Metatranscriptomics

Depletion of rRNA from extracted RNA was performed using QIAseq FastSelect −5S/16S/23S (Qiagen). Samples were incubated at 89°C for 8 minutes. Libraries for metatranscriptomics were prepared using the Illumina TruSeq® Stranded mRNA Library Prep Kit (Illumina) according to the manufacturer’s instructions. Shotgun metatranscriptomic sequencing of week 12 fecal samples was performed on an Illumina NextSeq 500/550 using the HighOutput Kit v2.5 (75 cycles).

Quality control and trimming were performed as described above for shotgun sequencing. Gene expression in the evolved population was calculated in Geneious (v 2022.2.1; https://www.geneious.com) by mapping reads to the reference genome of *E.coli* NC101 (PRJNA596436). Differential expression was calculated using DeSeq2^65^ as implemented in Geneious, using the parametric model and by assigning the conditions to “WT-” or “KO-evolved”.

### Measurement of the metabolic activity of bacteria

Biolog GENIII MicroPlates (Biolog, Hayward, CA, USA) was used to investigate the metabolism of evolved populations. Evolved populations stored in glycerol were inoculated onto the Inoculation Fluid-A (IF-A) provided by the manufacturer such that the OD_600_ was between 0.02-0.05. After adjusting the OD_600_, the inoculated IF-A was shaken well and distributed into a Biolog GENIII MicroPlate (100 μl in each well). The plate was then covered with a sterile Breathe-Easy membrane (Sigma-Aldrich, St. Louis, MO) and placed in a plate reader (SPARK, Tecan, Männedorf, Switzerland) at a preset temperature of 37 °C. The plate reader was then run with the following parameters: temperature: 37 °C, shaking: 250 rpm, OD_590_ measurement: every 15 min, total running time: 36 h. From the OD_590_ measures, area under the curve (AUC) was used as a proxy for metabolic activity. The Biolog assay was repeated for populations from fecal samples collected at weeks 1 and 12 from all 25 mice, with two replicates each.

### Ex vivo assay

The cecal content was collected from each mouse during dissection at the end of the second experiment. The amount of cecal content from each mouse was different based on the amount in the cecum at the time of dissection. Sterile PBS was added to the cecal content, and the volume was adjusted based on the amount of cecal content collected from each mouse (20 mg/ml). The protocol was adapted from Kitamoto et al. (2020) ^66^. The PBS-cecal content mixture was centrifuged twice (once at 500*g* for 5 min and again at 10000*g* for 5 min) and then filter-sterilized through a 0.2 μm filter. The *ex vivo* medium from each mouse was mixed well and 100 μl was spread onto an LB agar plate to ensure that they were sterile. *Ex vivo* media from all healthy mice were pooled, as were the *ex vivo* media from all inflamed mice, resulting in two *ex vivo* media. Populations from fecal samples collected at week 12 from a subset of mice (n = 5 KO and n = 5 WT mice) were inoculated into the *ex vivo* media and incubated in the TECAN Spark plate reader at a preset temperature of 37 °C. The plate reader was then run with the following parameters: temperature: 37 °C, shaking: 250 rpm, OD_600_ measurement: every 15 min, total running time: 36 h. From the OD_600_ measures, area under the curve (AUC) was used as a proxy for growth.

### Metabolomic evaluation of the fecal pellets

During the second experiment, samples for metabolomic investigation were collected once a week and snap-frozen in liquid nitrogen. Samples were stored at −70 °C and delivered at Helmholtz Center Munich for metabolomic investigation. A non-targeted metabolomics approach of mouse fecal samples was undertaken using NMR spectroscopy. To extract the aqueous metabolites, we homogenized 2-3 fecal pellets in 1 mL H_2_O using ceramic beads (NucleoSpin, Macherey–Nagel, Dueren, Germany) and a TissueLyser (Qiagen, Hilden, Germany), mixing the sample for 3 × 30 s at 4,500 rpm with a 10 s cooling break (< 0°C). Subsequently, the homogenate was centrifuged (13,000 rpm for 10 min at 4 °C), the supernatant evaporated using a SpeedVac, and the dried extract reconstituted in 200 µL NMR buffer (10% D_2_O, 100 mM phosphate buffer with 0.1% trimethylsilyl-tetradeuteropropionic acid (TSP), pH 7.4). Samples were transferred to 3 mm NMR tubes, and immediate NMR analysis was performed in a randomized order with a Bruker 800 MHz spectrometer operating at 800.35 MHz equipped with a Bruker SampleJet for sample cooling (283 K) and a QCI-cryogenic probe. A standard one-dimensional pulse sequence (noesygppr1d) provides an overview of all molecules. The acquisition parameters were as follows: water suppression irradiation during recycle delay (2 s), mixing time of 200 ms, 90 °pulse of 12.5 µs. We collected 512 scans of 64 K data points with a spectral width of 12 ppm. The software TopSpin 3.6 (Bruker BioSpin, Ettlingen, Germany) was used for processing, i.e., Fourier transformation, manual phasing, baseline correction, and calibration to TSP (δ 0.00). Data were imported into Matlab software R2011b (Mathworks, Natick, MA, USA) and further processed, i.e., the water region was removed, baseline adjusted ^67^ and spectra normalized ^68^. Relative quantification of metabolites was performed using the peak heights of selected peaks and compounds identified as described in our published workflow ^69^.

### Shortest path analysis

For the shortest path analysis, we used the metabolic model reconstructed using the AGORA2 resource for the human microbiome ^32^. The model was translated into a graph with metabolites as nodes, and the reactions converting metabolites into one another as edges. The shortest path analysis was performed using the Dijkstra algorithm implemented in the igraph package (1.3.5) of R (4.2.2) ^70,71^. To avoid shortcuts through the network using cofactors as intermediate compounds, the edges were weighted by the sum of degree of the nodes it connected, as suggested by Faust et al. ^72^. For each compound from the *in vitro* screen, we calculated the set of shortest paths to all other metabolites in the model. Each value in the set was the average shortest path of the five shortest paths between two metabolites, which was performed to account for uncertainties in pathway calculation based only on the network properties. For our analysis, we only considered pathways between compounds from the two datasets, which were among the shortest 5% of pathways in a pathway set.

### Prediction of possible conversions between metabolites

We wanted to understand whether the altered metabolic capabilities observed in the Biolog plates or the changed environmental conditions observed in the metabolomic data, together with the metabolism of *E. coli* could contribute to the observed changes in the metabolomic data between KO and WT mice. To this end, we performed flux variable analysis (FVA) on the metabolic model for E. coli NC101 (retrieved from AGORA2) ^32^. We defined every significantly detected metabolite in the Biolog experiment or the metabolomic data set as a possible source metabolites, while the metabolites from the metabolomic data set were defined as target metabolites. Metabolites found in the metabolomics are those known to be present *in vivo*. Compounds found to be significantly differentially metabolized in the Biolog assay are those that we know the evolved bacteria are capable of metabolizing. By using the sum of these as the source and the detected metabolites as the target, we aimed to capture as many of the possible conversions as possible to help explain the composition of the *in vivo* environment and the role of bacterial metabolism in shaping it. For the simulation background, we employed a minimal medium (Table S13), removed D-Glucose, and adjusted the oxygen level according to the conditions tested (anoxic = 0 mmol h⁻¹ gDW⁻¹, microaerobic = 1 mmol h⁻¹ gDW⁻¹, aerobic = 10 mmol h⁻¹ gDW⁻¹). For the simulation, we added each source metabolite individually to the growth medium (100 mmol h⁻¹ gDW⁻¹) and calculated the maximum production rates of the target metabolites, assuming that at least 50% of maximum growth rates were achieved. We considered a maximum flux of >1e⁻⁶ as the possible production of the target metabolite from the source metabolite. FVA was performed in cobrapy ^73^ and analyses were performed using the data.table and ggplot2 packages for R.

### Statistical investigation

Statistical analyses were performed using packages in RStudio 2023.03.0+386. For PERMANOVA analyses, adonis was used ^74^ on Bray-Curtis distances of the data ^75^ and mixomics was used for PLS-DA ^76^. AUCs were calculated using desctools ^77^.

## Acknowledgments

We are grateful to Balfour Sartor for providing the NC101 *E. coli* strain, the MPI-Plön mouse team for support with animal experiments, Britt Marie Hermes for support in manuscript preparation, and Wael Elhenawy and Isabel Gordo for their helpful and constructive discussions. The authors acknowledge the use of ChatGPT for the purpose of supporting the drafting of material for the introduction section.

## Data availability statement

*Escherichia coli* NC101 reference genome is available under Bioproject number PRJNA596436. Shotgun metagenomic sequencing of the inocula and evolved populations and metatranscriptomic of the populations at week 12 are available with Bioproject number PRJNA1012288.

## Author contributions

MV, DU, and JB conceptualized the study; RU, NAA, MV, SH, MAG, OV, SK and DU performed experiments; RU, NAA, MV, SH, FT, JH, and PR analyzed the data; JT and CK performed the metabolic modeling; RU, NAA, and AD performed the statistical analyses. RU, NAA, DU, and JB wrote the first draft of the manuscript. All coauthors revised the manuscript and agreed to its publication.

