## Supplementary figures for "Evolution of *E. coli* in a mouse model of inflammatory bowel disease leads to a disease-specific bacterial genotype and trade-offs with clinical relevance"

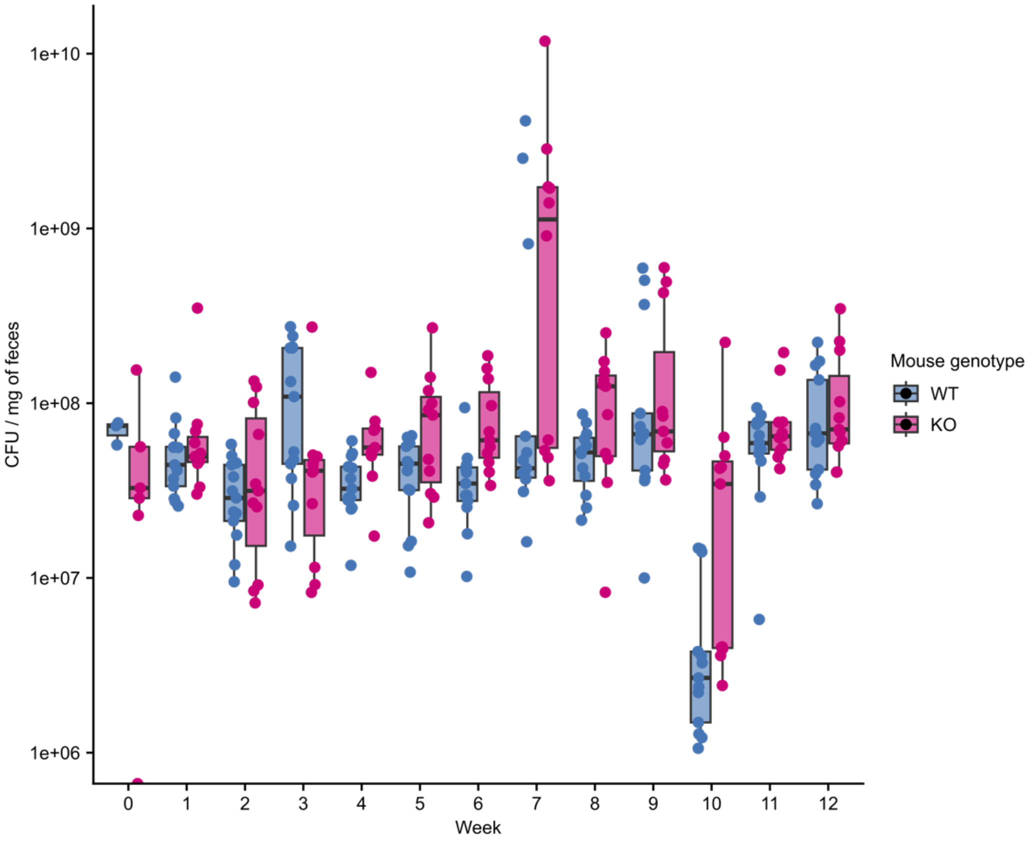


Figure S1: Fecal bacterial load is stable during the experiment
Bacterial counts (CFU / mg of feces) at different timepoints during the experiment. Each dot represents bacterial load of feces from independent mice at each time point. Pairwise comparisons of bacterial loads of feces between the two mouse genotypes are reported in Table S5. Differences were considered statistically significant at Benjamini-Hochberg-corrected $P < 0.05$.

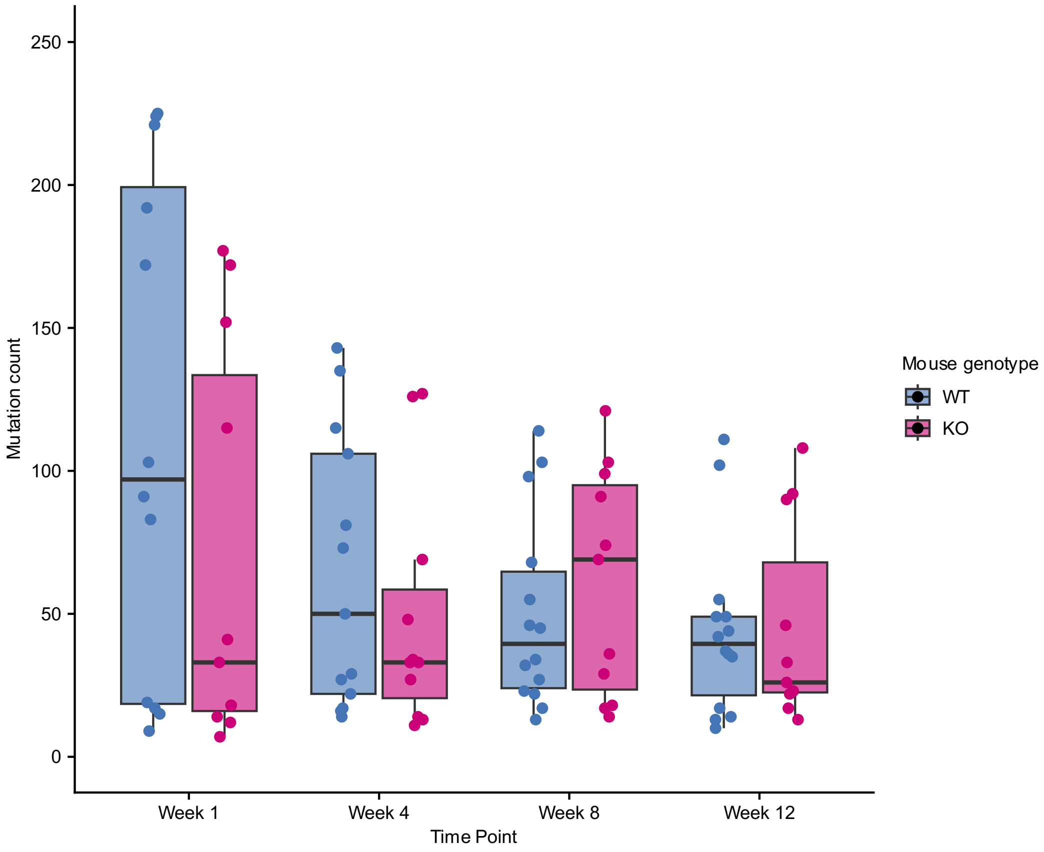


Figure S2: number of *de-novo* mutations in populations evolved in WT and KO mice at week 1, 4, 8 and 12

*De-novo* mutation counts in populations evolved in WT and KO mice at week 1, 4, 8 and 12. Each dot represents the populations of one mouse. Pairwise comparisons of mutation counts between populations evolved in the two mouse genotypes are reported in Table S7. Differences were considered statistically significant at Benjamini-Hochberg-corrected $P < 0.05$.

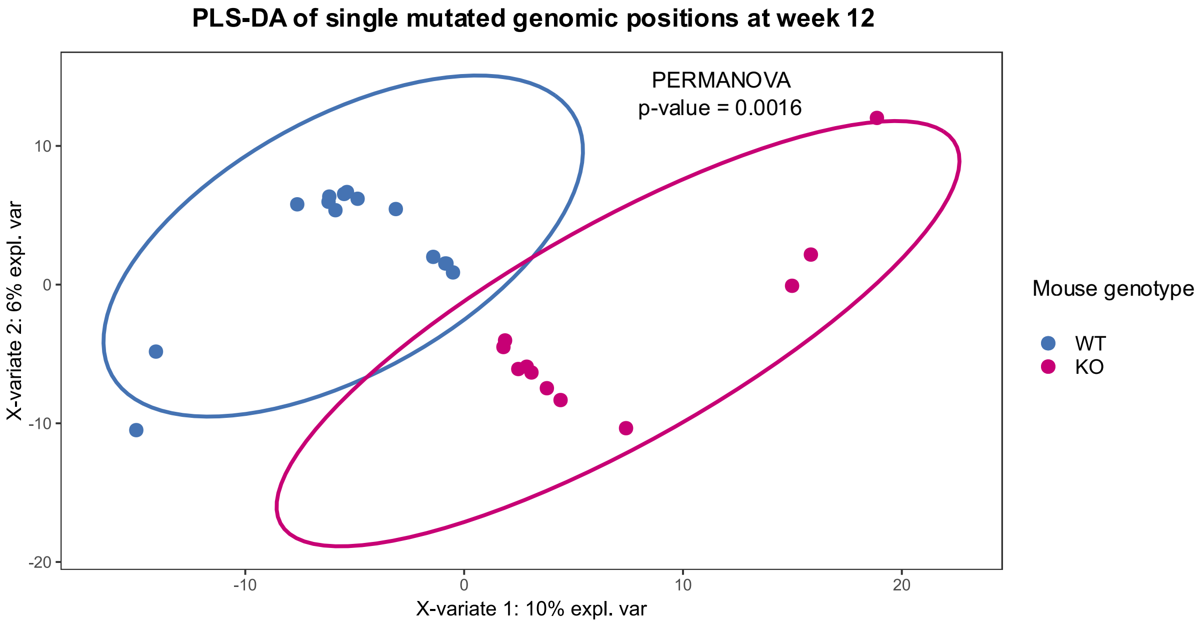


Figure S3: *De-novo* mutations at individual nucleotides differed significantly between bacteria from WT and KO mice. Similar to mutated loci, mutated positions also showed a discrete clustering of bacterial populations from WT and KO mice upon performing PLS-DA. PERMANOVA $P < 0.05$.

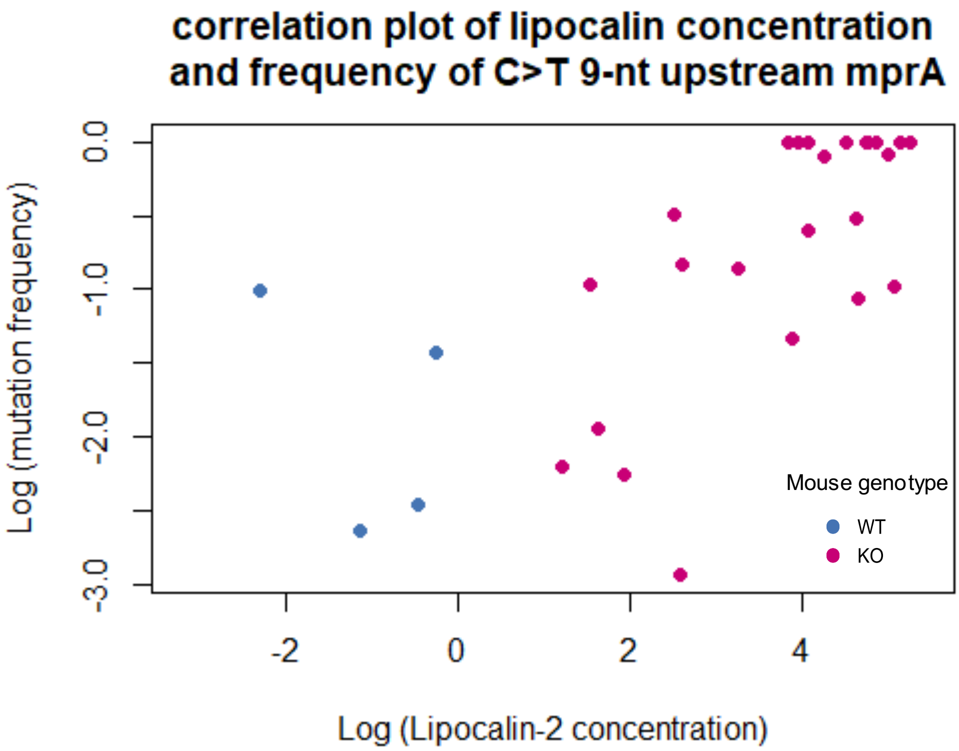


Figure S4: Frequency of the candidate mutation (SNP upstream of *mprA*) in the bacterial populations in relation to inflammation, as measured by fecal lipocalin-2 at all sequenced timepoints

Log-lipocalin-2 levels and log-frequency of the candidate mutation at weeks 0, 1, 4, 8 and 12 are plotted. The increase in frequency of the C>T mutation strongly coincides with the increase in inflammation as measured by lipocalin-2 levels, both in KO mice alone, and all mice are included (Fig 2C, Fig S4; Spearman's rank correlation rho $P$ = 0.00005 for KO mice alone and $P$ = 3.005e-11 for all mice).

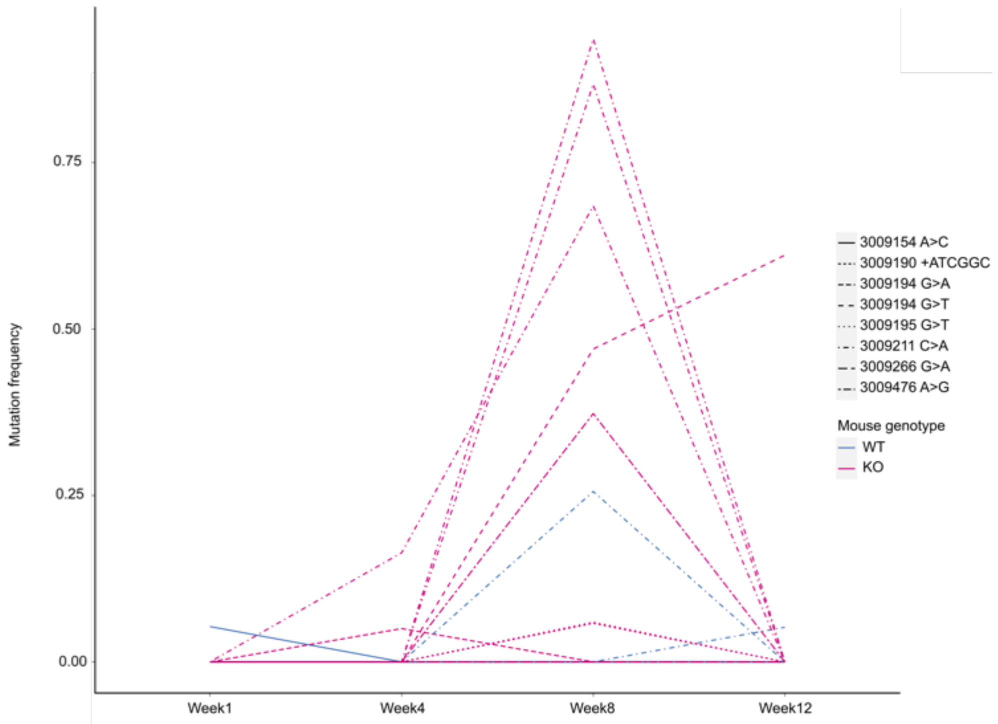


Figure S5: The genetic locus between hypothetical protein and *mprA* is a mutational hotspot Apart from the candidate mutation described in the main text (C>T change at position 3009211), other mutations were also observed at different points in the experiment across mice. The graph shows the changes in frequencies of other mutations in the same region.

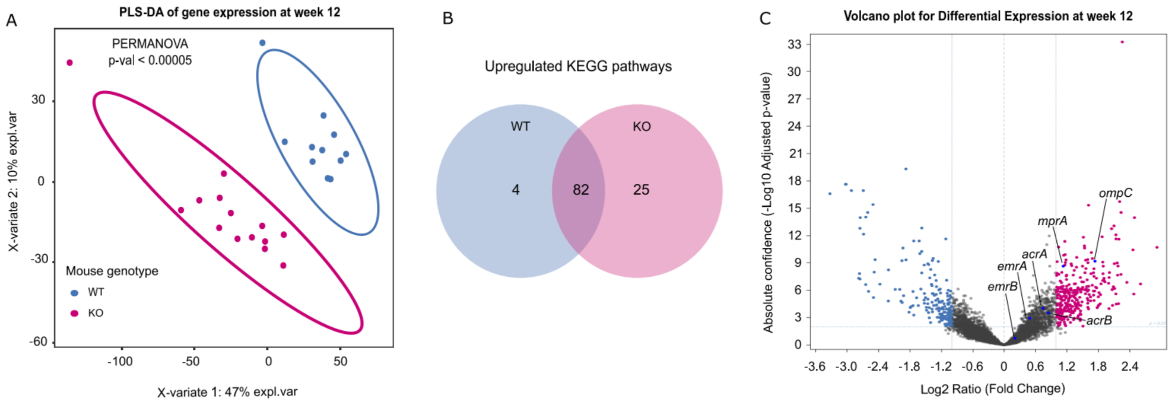


Figure S6: Gene expression profiles of *E.coli* populations in the healthy and inflamed gut
A Partial least squares-discriminant analysis of the expressed genes (using RPKM as a measure of expression) in the evolved populations of *E. coli* NC101 at week 12 in WT and KO mice. Differences were considered statistically significant at a PERMANOVA $P < 0.05$; B Differentially expressed genes were classified based on KEGG pathways; C Volcano plot showing statistical significance (adjusted P-value) versus magnitude of change (Log2 Ratio [fold change]) of up-regulated genes in populations at week 12 in KO (blue) and WT (pink) mice. The genes belonging to the *mprA* regulon are highlighted in dark blue in the plot.

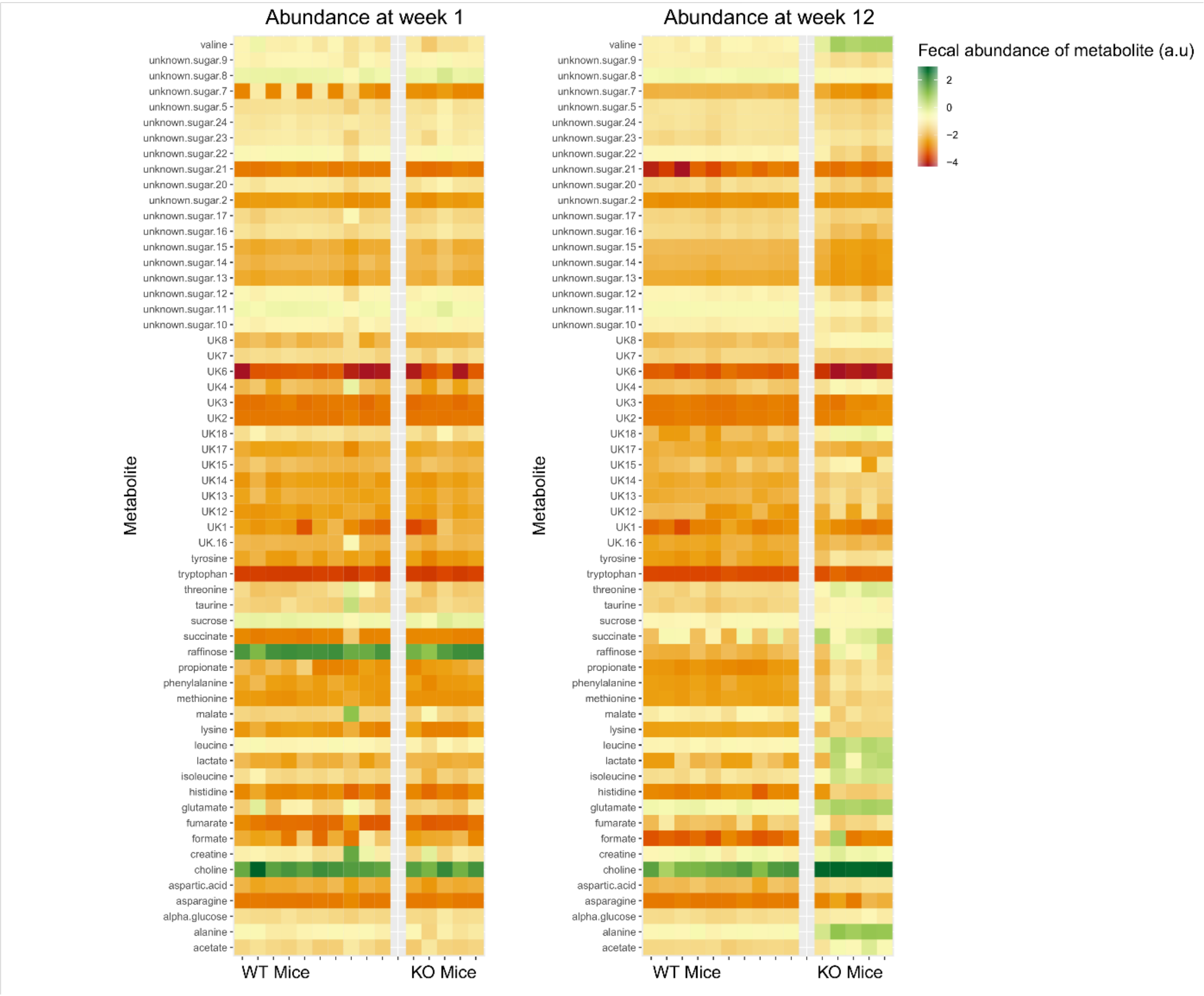


Fig S7: Heatmap showing the log of fecal abundance (arbitrary unit; a.u.) of the identified compounds from metabolomics in samples from weeks 1 and 12. Each column represents a mouse from Experiment 2 (genotype is indicated on the X axis) and each row indicates the detected compound.

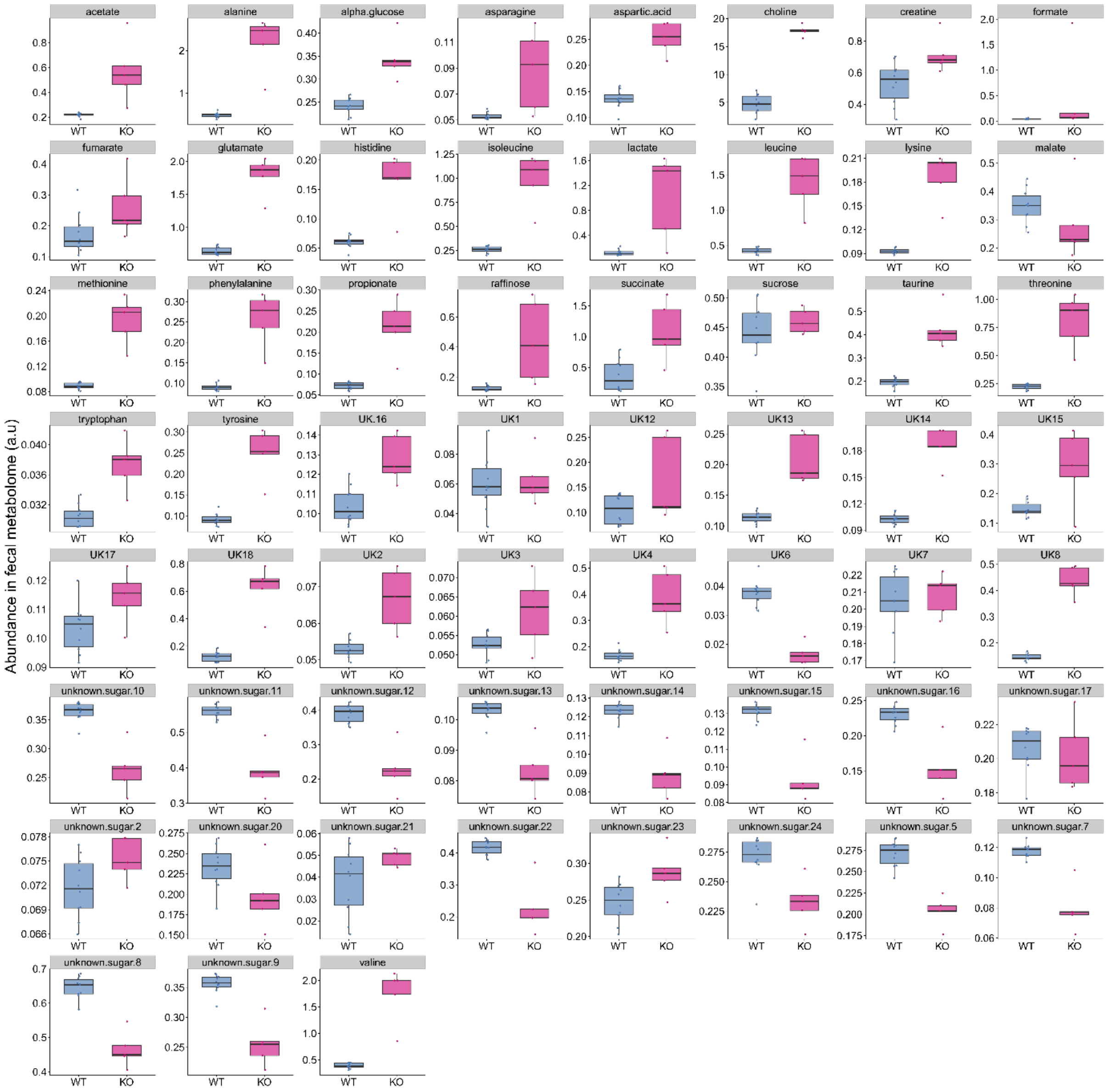


Figure S8: The metabolites in which fecal abundance was significantly different between the WT and KO mice at week 12 are represented in each box. Fecal abundance is shown on the Y axis (arbitrary unit; a.u.) and the mouse genotype on the X axis. For all shown metabolites, the difference between WT and KO mice is statistically significant ($P < 0.05$).

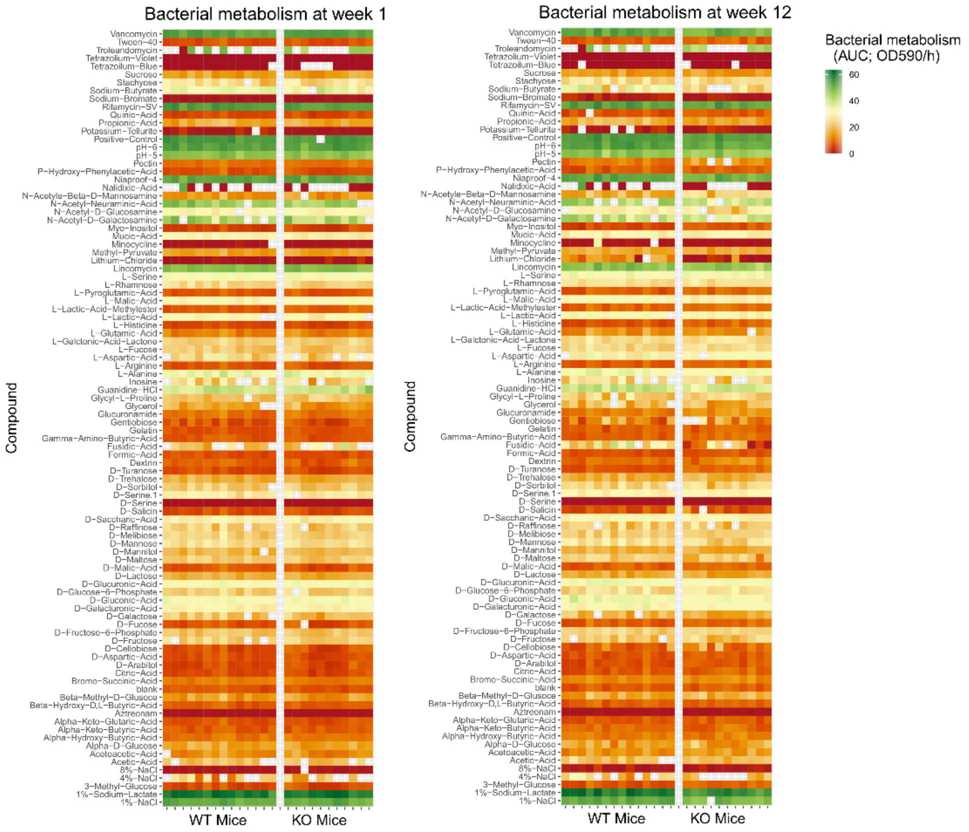


Fig S9: Heatmap showing the metabolic activity (AUC of metabolic activity measured in OD590/hours) in the presence of the different conditions on the BIOLOG GENIII plate. Data of bacterial populations from week 1 and week 12 are shown in the two panels. Each column represents a mouse (genotype is indicated on the X axis) and each row indicates the condition on the BIOLOG GENIII plate. Blank cells indicate the data was not considered in the analysis because the measurements from the two replicates were very different (greater than 10 OD590/h).

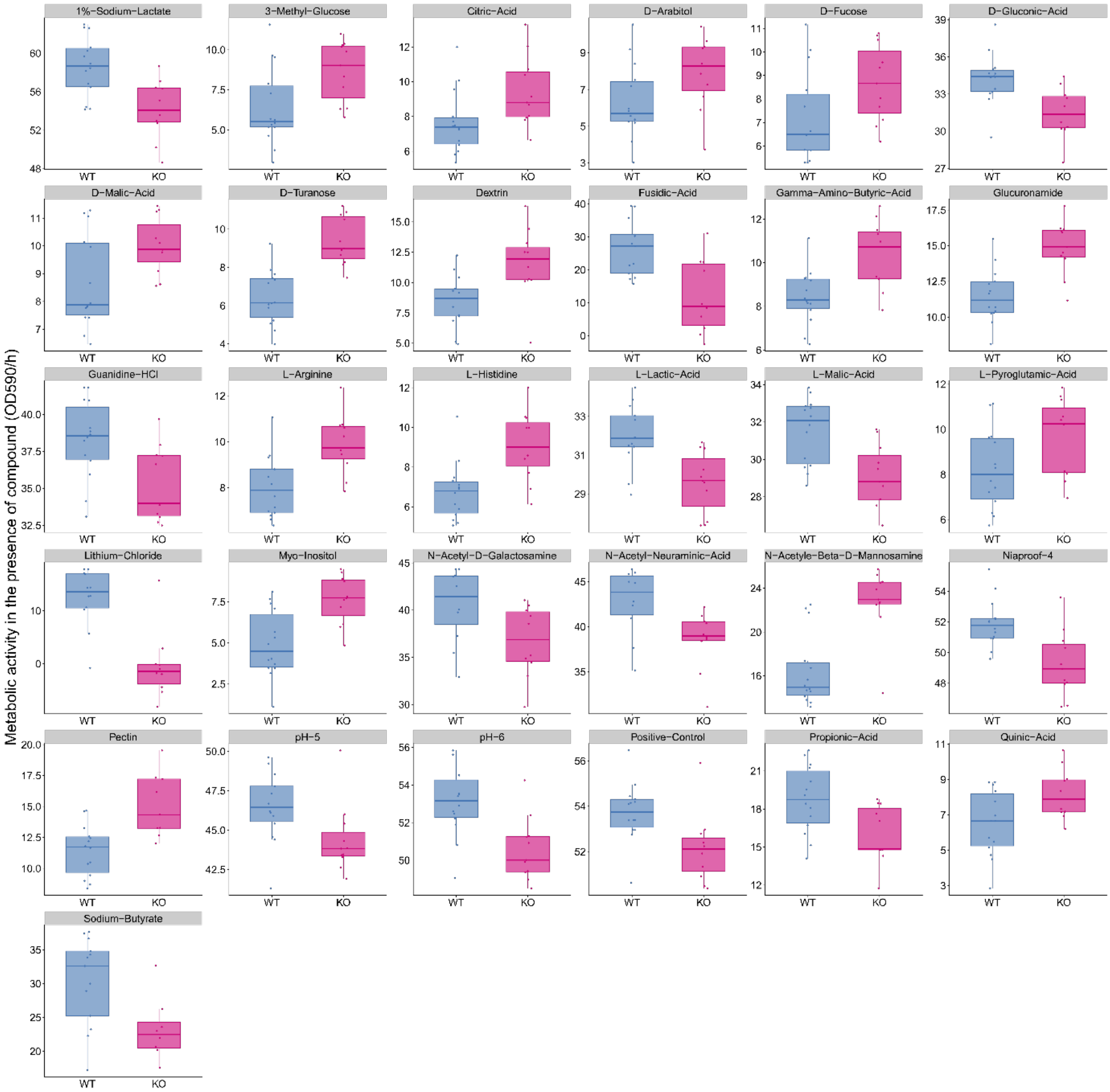


Figure S10: The compounds/conditions in which the bacteria from WT and KO mice at week 12 show significantly different metabolic activity are represented in each box. Metabolic activity is shown on Y axis, expressed as AUC (O.D.590/hours). and the mouse genotype is shown on X axis. For all compounds/conditions shown, the metabolic activity is significantly different between WT and KO mice ($P < 0.05$).

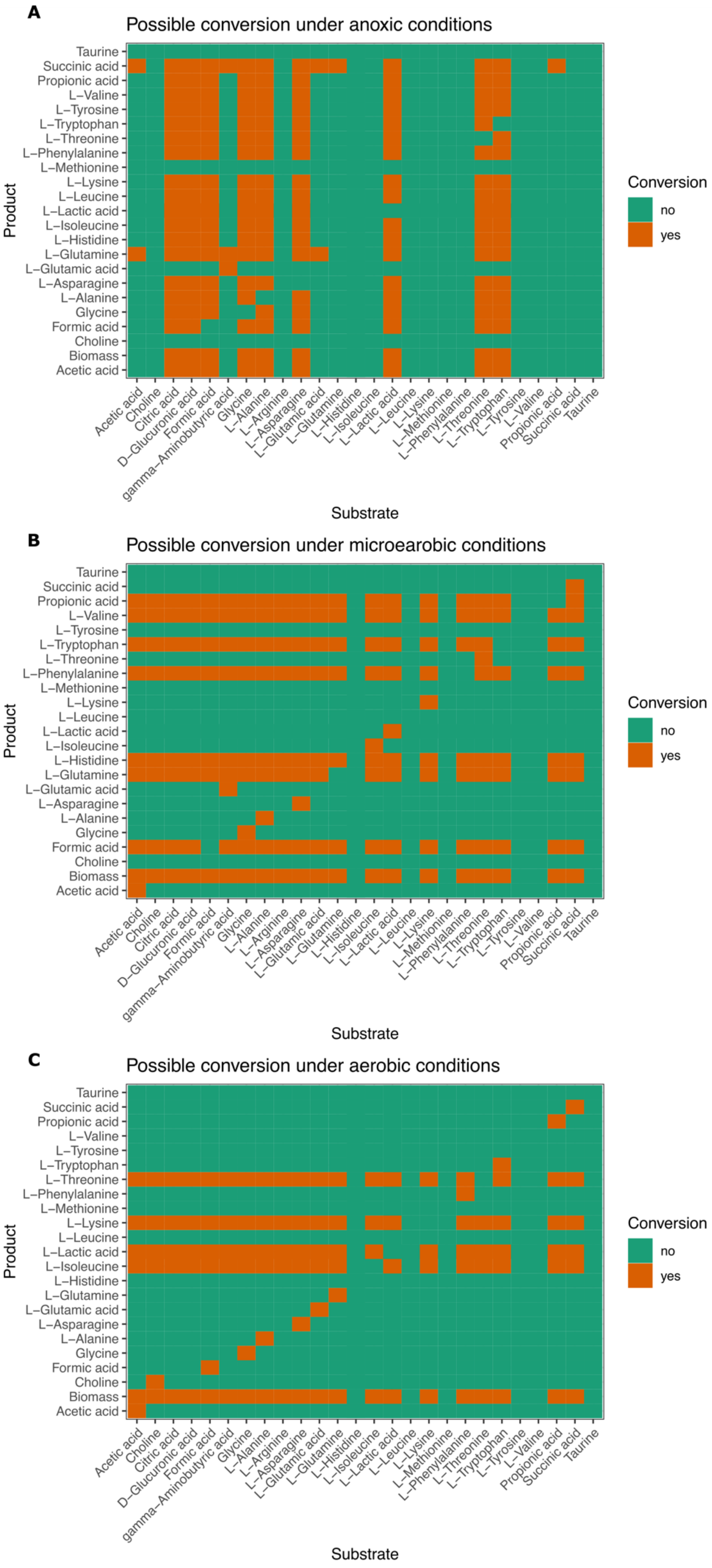
A
Possible conversion under anoxic conditions
Product
Taurine
Succinic acid
Propionic acid
L−Valine
L−Tyrosine
L−Tryptophan
L−Threonine
L−Phenylalanine
L−Methionine
L−Lysine
L−Leucine
L−Lactic acid
L−Isoleucine
L−Histidine
L−Glutamine
L−Glutamic acid
L−Asparagine
L−Alanine
Glycine
Formic acid
Choline
Biomass
Acetic acid
Substrate
Acetic acid
Choline
Citric acid
D−Glucuronic acid
Formic acid
gamma−Aminobutyric acid
Glycine
L−Alanine
L−Arginine
L−Asparagine
L−Glutamic acid
L−Glutamine
L−Histidine
L−Isoleucine
L−Lactic acid
L−Leucine
L−Lysine
L−Methionine
L−Phenylalanine
L−Threonine
L−Tryptophan
L−Tyrosine
L−Valine
Propionic acid
Succinic acid
Taurine
Conversion
no
yes
B
Possible conversion under microearobic conditions
Product
Substrate
Conversion
no
yes
C
Possible conversion under aerobic conditions
Product
Substrate
Conversion
no
yes

Fig S11: Flux variability analysis on the metabolic model of *E. coli* NC101. The graphs show *in silico* predictions of whether *E. coli* NC101 can produce certain compounds (Y axis) from other compounds (X axis). Each compound on the X axis was assumed to be added to a minimal media for this analysis. The row “Biomass” on the Y axis indicates whether *E. coli* NC101 can utilize the compounds on the X axis for growth. Panels **A**, **B**, and **C** show the results of the flux variability analysis in anoxic, micro-aerobic, and aerobic conditions, respectively.

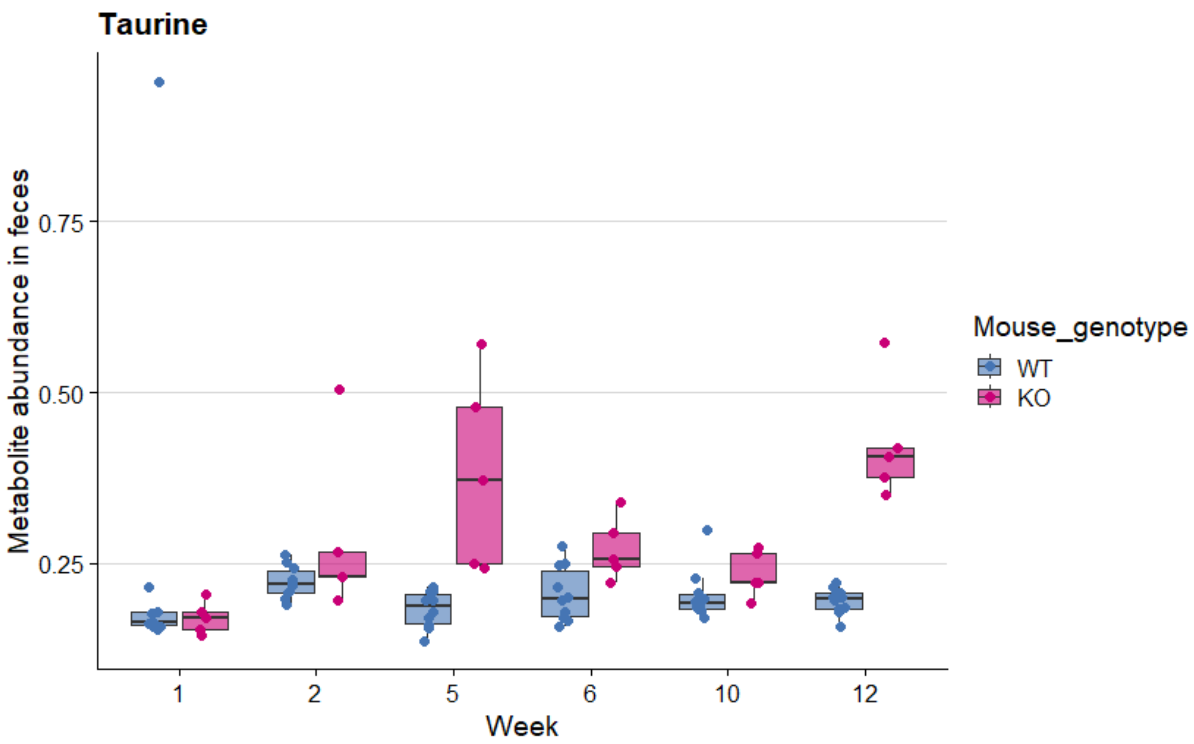


Fig S12: Change in fecal abundance of taurine at different points during the experiment in WT and KO mice.

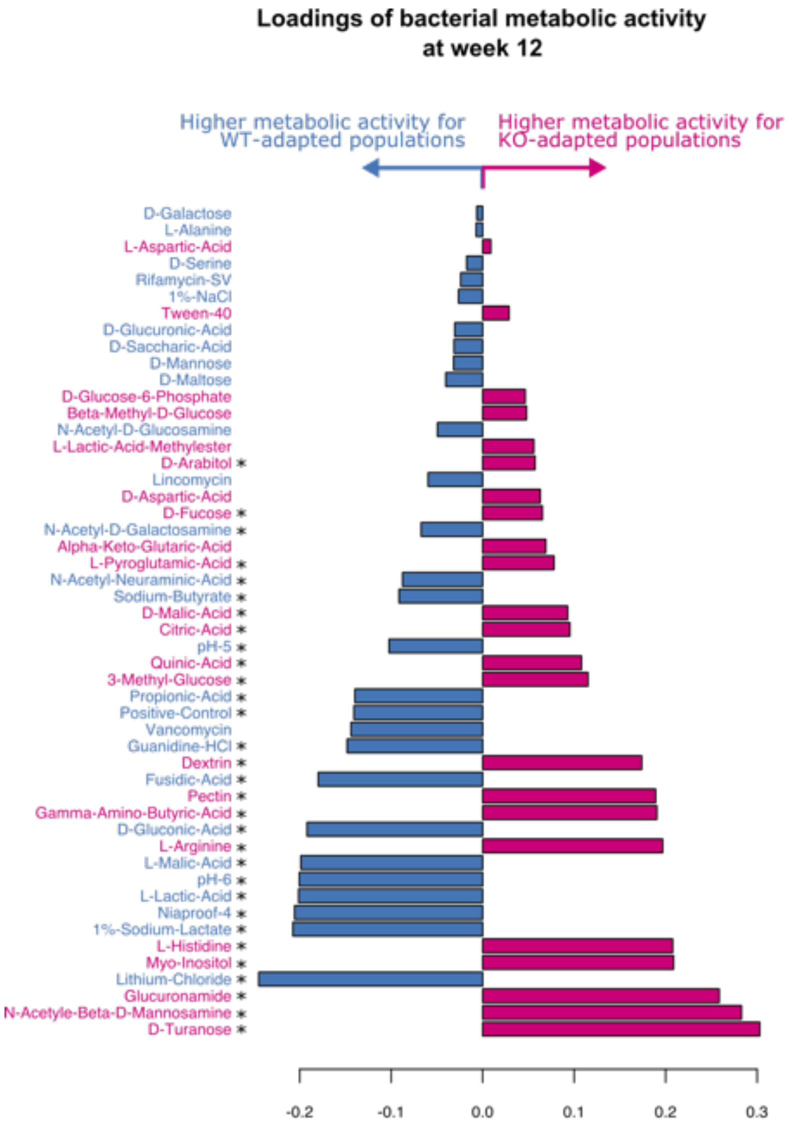


Figure S13: Loadings of the compounds in the *in vitro* metabolic screen are shown, i.e., compounds that contributed to the difference in metabolic activity between the two bacteria are shown. The compounds are arranged based on the weight of each compound's contribution to the observed difference, with the compound with the highest weight at the bottom. Significance stars indicate whether the compound induced significantly different activity in the bacteria, based on univariate analysis. *, Kruskal-Wallis H test at Benjamini-Hochberg corrected $P < 0.05$
